# Beyond genetic indicators: how reproductive mode and hybridization challenge freshwater mussel conservation

**DOI:** 10.1101/2025.03.28.645956

**Authors:** Ellika Faust, Julie Conrads, Marco Giulio, Claudio Ciofi, Chiara Natali, Philine G.D. Feulner, Alexandra A.-T. Weber

**Affiliations:** Department of Aquatic Ecology, Swiss Federal Institute of Aquatic Science and Technology (Eawag), Dübendorf, Switzerland; Department of Fish Ecology and Evolution, Swiss Federal Institute of Aquatic Science and Technology (Eawag), Kastanienbaum, Switzerland; Department of Biology, University of Florence, Florence, Italy; Institute of Ecology and Evolution, University of Bern, Bern, Switzerland

## Abstract

Genetic diversity is a fundamental component of biodiversity that should be monitored and preserved. Unionid freshwater mussels provide essential ecosystem services but are among the most imperilled aquatic organisms worldwide. Here, we assess key genetic Essential Biodiversity Variables (EBVs)—genetic diversity, genetic differentiation, inbreeding, and effective population size—in all *Anodonta* species in Switzerland. After generating draft genomes for *Anodonta cygnea* and *Anodonta anatina,* we performed whole-genome resequencing of 421 individuals (243 *A. anatina*, 151 *A. cygnea*, 17 *A. exulcerata* and 10 *Anodonta* sp.) from 31 populations. While *A. anatina* populations followed a metapopulation structure shaped by catchment areas, genetic diversity correlated positively with waterbody size, suggesting greater vulnerability in small ponds compared with large lakes. Inbreeding levels were low, however effective population sizes were consistently below 100, indicating serious extinction risks. We also detected hybridization between *A. cygnea* and *A. exulcerata*, indicating genomic permeability between these species. Furthermore, genomic data suggested facultative selfing in *A. cygnea*, leading to a marked reduction in genetic diversity, increased population structure and inbreeding, and a decline in effective population size compared to the outcrossing *A. anatina*, emphasizing that *A. cygnea* faces a particularly high risk of extinction due to its reproductive strategy. Our study reveals the vulnerability of freshwater mussels and emphasizes the need for genetic indicators to reflect species-specific reproductive strategies. More broadly, it calls for conservation policies to integrate genetic monitoring and consider reproductive modes to tailor conservation efforts and better assess extinction risks.

## Introduction

The Anthropocene is characterised by a global biodiversity crisis, with unprecedented species extinction caused by human activities (Johnson et al., 2017). The importance of intraspecific genetic diversity has been recognized in the global biodiversity framework (GBF) of the convention on biological diversity (CBD), which acknowledges that genetic diversity is key for population persistence and adaptation to changing environments and extreme events. Endangered populations and species are typically characterised by low genetic diversity and are threatened by genetic erosion that can reduce fitness and ultimately contribute to extinction (Kardos, 2021; Kardos et al., 2021). Worryingly, comparative meta-analyses have highlighted that genetic diversity is decreasing globally (Leigh et al., 2019; Shaw et al., 2025). Hence it is imperative that conservation actions consider and integrate this fundamental level of biodiversity (O’Brien et al., 2022, 2025). Essential Biodiversity Variables (EBVs) are a standardized set of measurements designed to monitor biodiversity changes across time and space. The genetic composition EBVs focus on the genetic diversity within and among populations of species, specifically genetic diversity, genetic differentiation, inbreeding, and effective population size (Hoban et al., 2022; Paez et al., 2022). These estimates can help to identify conservation units, assess population connectivity, and biodiversity loss (Hohenlohe et al., 2021). Beyond EBVs, genomic data can also reconstruct demographic history, detect hybridization, evaluate a population’s adaptive potential (Hohenlohe et al., 2021) and uncover cryptic species (Bickford et al., 2007; Struck et al., 2018). Clear species definitions are crucial in conservation, as actions largely occur at the species level (e.g., IUCN Red List). Undescribed cryptic species risk being overlooked, potentially leading to underestimated vulnerability or unnoticed extinctions (Carmi et al., 2016).

The global decline in biodiversity is especially pronounced in freshwater ecosystems, where 85% of assessed biodiversity has been declining since the 1970s—significantly higher than the 56% and 69% observed in marine and terrestrial ecosystems, respectively (WWF, 2024). This marked decline is driven by two key factors (Dudgeon et al., 2006; Sayer et al., 2025). First, freshwater habitats are inherently vulnerable due to their fragmented nature, limited connectivity, and restricted dispersal via river networks. Second, freshwater ecosystems face intense anthropogenic pressure, stemming from the direct use of water resources for purposes such as wastewater and drinking water plants, dams and hydropower, fishing, irrigation, channelization, and pollution from land use. Freshwater mussels (Bivalvia: Unionida) are considered keystone species, integral to the health of these ecosystems, providing critical services such as nutrient cycling and water purification (Vaughn, 2018). Globally, nearly half of the 456 species evaluated by the IUCN are classified as threatened or near-threatened (Böhm et al., 2021). This decline is driven by several factors, including habitat destruction and modification, invasive species, pollution and climate change (Lopes-Lima et al., 2014, 2017). Their life-history traits make them particularly vulnerable to environmental disturbances: they exhibit high longevity, reach sexual maturity late, and have a complex life cycle that includes an obligatory parasitic larval stage on a host fish (Lopes-Lima et al., 2017). While some species are generalists regarding host fish species, others have strong host specificity, making them particularly vulnerable when host fish populations are also declining (Modesto et al., 2018). Importantly, this larval parasitic stage is the primary mechanism of mussel dispersal and colonization of new habitats (Modesto et al., 2018).

While 25 species of freshwater mussels have been described in Europe, the true diversity of these mussels is likely underestimated due to the reliance on shell morphology for species identification, which is highly variable. This has led to historical splitting and lumping of species (Haas, 1969; Graf, 2010). However, recent molecular studies have shown that some splits were justified, such as the resurrection of *Anodonta exulcerata* as a distinct species from *Anodonta cygnea* (Froufe et al., 2017; Riccardi et al., 2020). Additionally, the *Unio crassus* species complex has been recently completely re-evaluated, revealing 12 biological species (Lopes-Lima et al., 2024). Molecular studies have also revealed that the widespread *A. anatina* likely represents a complex of cryptic species, with five divergent mitochondrial lineages and at least three evolutionary significant units, which may represent cryptic species (Froufe et al., 2014, 2017; Lopes-Lima et al., 2016; Riccardi et al., 2020; Lyubas et al., 2023). Despite extensive mitochondrial data, few studies have analysed nuclear data, and then only a limited number of markers (Lopes-Lima et al., 2016). Hence, there is to date no study that used whole-genome data for species delimitation and for the assessment of genetic EBVs in freshwater mussels.

This study focuses on Switzerland, as conservation policies, jurisdiction, and biodiversity reporting are implemented at the national level. Despite its small geographic extent, the country plays a crucial role in the biogeography of freshwater biodiversity and is often referred to as the “water castle of Europe.” The Alps give rise to several major river catchments, including the Rhine, Rhône, and Ticino, as well as key sub-catchments such as the Limmat, Reuss, and Aare, all of which ultimately flow into the Rhine. Given this complex hydrological network, strong genetic differentiation over short geographic distances is expected. In Switzerland, in addition to *A. anatina,* two other *Anodonta* species are present: *A. exulcerata* and *A. cygnea* (Pfarrer et al., 2022). While *A. anatina* is typically gonochoric (although occasional hermaphroditism has been reported (Hinzmann et al., 2013)), *A. cygnea* is known to be predominantly hermaphroditic (Lima et al., 2012), with laboratory studies reporting instances of self-fertilization (Bloomer, 1940, 1943). However, the prevalence of self-fertilization in natural populations remains unclear. Despite their coexistence in Switzerland, *A. anatina* and *A. cygnea* are not closely related, with an estimated divergence time of around 25 million years (Lopes-Lima et al., 2021, 2024). Both species are considered host generalists; however, some species-specific preferences have been described (Huber and Geist, 2017, 2019). The aim of this study was to assess genetic EBVs using high-quality genomic data for all *Anodonta* species present in Switzerland, in alignment with the reporting requirements of the CBD. In addition to estimating EBVs, our genomic analyses revealed cryptic diversity, evidence of hybridization, and insights into genetic consequences of a shift in reproductive strategy.

## Methods

### Population genomics of *Anodonta* mussels

Methods related to *A. cygnea* and *A. anatina* draft genome generation are available in the supplementary methods. Collection permits for population sampling were obtained in advance to the relevant cantonal authorities of nature protection. *Anodonta* mussels were sampled across 31 sampling sites in Switzerland from May to September 2023 (Table 1). The sites were selected based on mussel observations reported in the Swiss Center for the Cartography of the Fauna (www.infofauna.ch), with the goal of covering all six major river catchments and sub-catchments: Aare, Limmat, Reuss, Rhine, Rhône, and Ticino. In total, 421 *Anodonta* specimens were collected by wading, snorkelling, or SCUBA diving, with a maximum of 25 individuals per location (Table 1; Table S1). After morphological examination, a small foot biopsy was sampled non-lethally and immediately preserved in 100% ethanol. After tissue sampling, all mussels were placed back in their original sampling locations. Due to the morphological overlap among *Anodonta* species commonly occurring in sympatry, molecular species identification was performed using COI barcoding as described in the supplementary methods. DNA extracted from each individual was used for individual library preparation with the PCR-free DNA Prep Kit (Illumina). All libraries were sequenced with 150 bp paired-end reads on a single 25B flow cell using a Novaseq X Plus sequencer at the Functional Genomics Center Zürich, Switzerland.

**Table 1:**
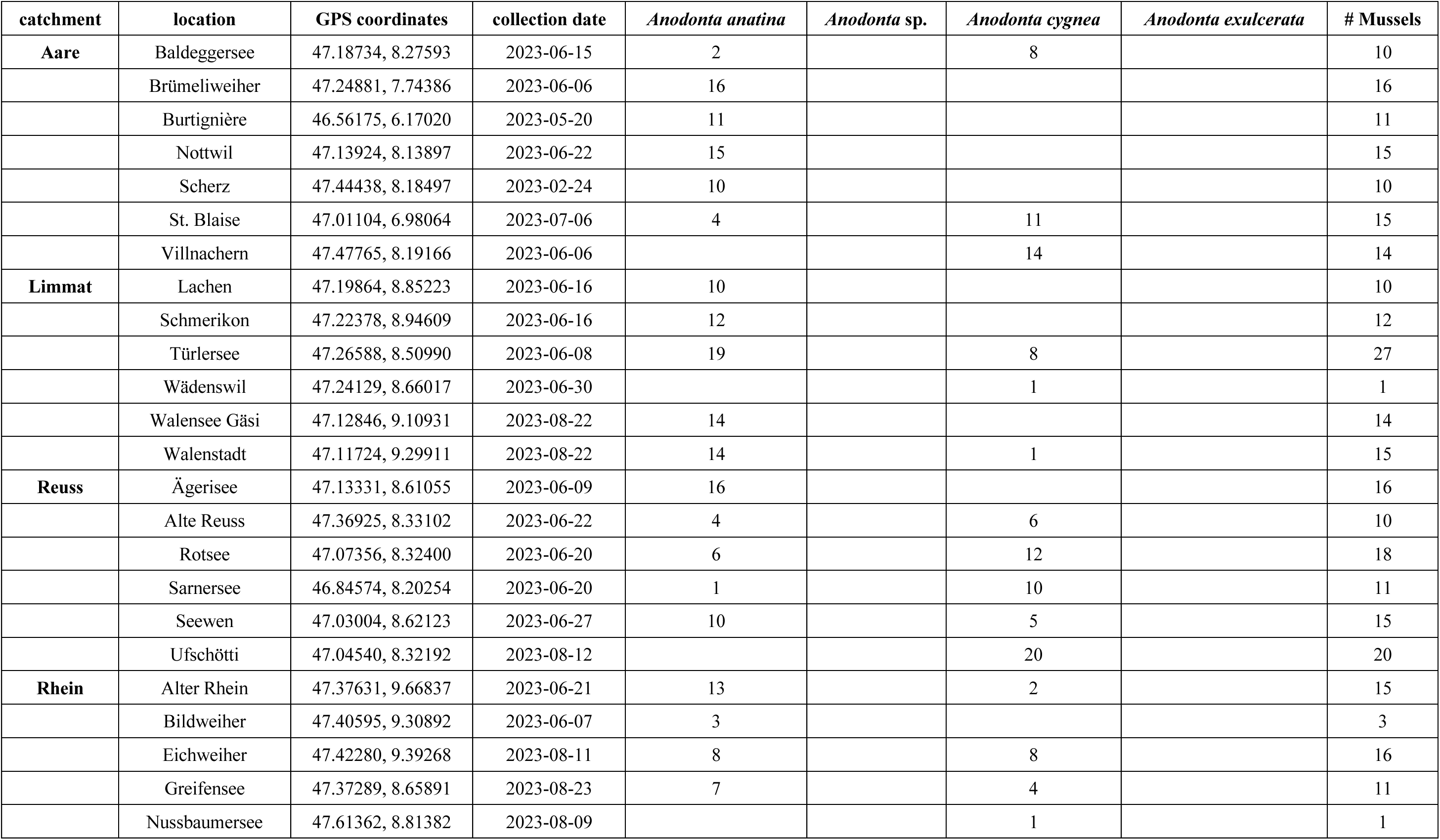
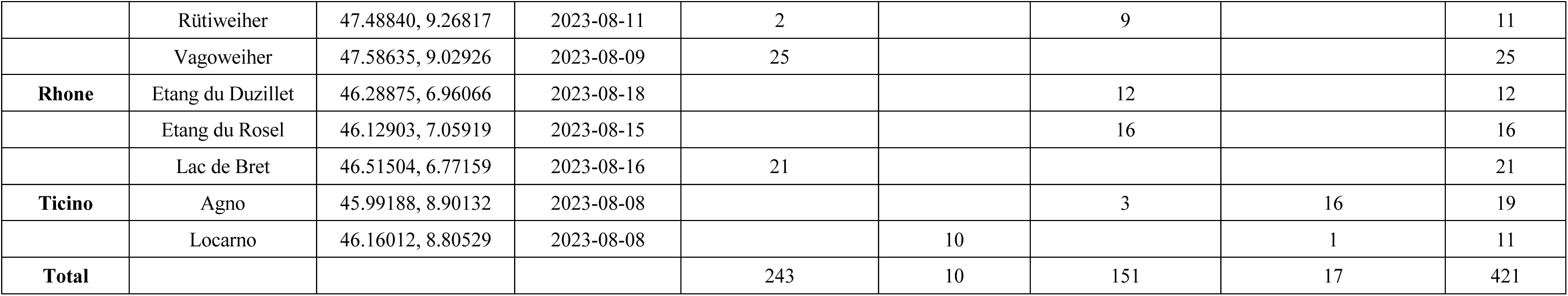
Sampling locations, dates and number of individual mussels collected during fieldwork.

#### Data processing and analyses

Following an initial quality check with FastQC 0.12.1 (Andrews, 2010), sequencing reads from all individuals were first mapped to the *A. anatina* genome assembly using BWA-MEM2 (Vasimuddin et al., 2019) to analyse all species together (dataset 1: all species). Reads with a mapping quality score below 20 were removed using Sambamba 1.0.1 (Tarasov et al., 2015), and optical/PCR duplicates were filtered out using Picard MarkDuplicates 3.3.0. (https://broadinstitute.github.io/picard/). Genotype calling was performed with bcftools 1.20 (Danecek et al., 2011), followed by filtering with bcftools, vcftools 0.1.16 (Danecek et al., 2011), and plink PLINK 1.9 (Purcell et al., 2007) (see Table S2 for filtering parameters). Linkage disequilibrium (LD) pruning was performed using PCAone (Bercovich et al., 2025), 2025) in 100 kb windows with a r² threshold of 0.1 and k=3. PCAone calculates an adjusted measure of LD by using the top inferred principal components to correct for population structure and admixture.

To improve genomic inference, resequencing data were mapped to the least divergent or species-specific reference genome, resulting in two datasets (*A. anatina* and *A. cygnea*+*A. exulcerata*). Mussels were first classified as *A. anatina*, *A. cygnea*, or *A. exulcerata* based on their COI barcode. This classification determined which sequences were mapped to the *A. anatina* or *A. cygnea* genome. For *A. exulcerata* individuals, the genome of *A. cygnea* - its closest sister species - was used for mapping. Sequences were processed the same way as for dataset 1 described above (Table S2). After filtering, the average sequencing depth ranged from 2.6x to 8.8x for the *A. anatina* dataset, while it ranged from 2.5x to 10.4x for the *A. cygnea*+*A. exulcerata* dataset. The final three filtered datasets contained the following number of single nucleotide polymorphisms (SNPs): all species —400 individuals and 26,704,704 SNPs; *A. anatina*—243 individuals and 27,996,116 SNPs; and the *A. cygnea*+*A. exulcerata* dataset— 155 individuals and 22,403,827 SNPs.

#### Data analyses

PCA was performed on the LD-filtered data in PLINK 1.9 (Purcell et al., 2007). Admixture proportions were estimated using ADMIXTURE 1.3 (Alexander et al., 2009). For the admixture analysis of *A. anatina,* the LD-pruned data was downsampled to 2 million SNPs for computational efficiency. Pairwise *F*_ST_ values were computed in PLINK 2 (Chang et al., 2015) using the Hudson method (Hudson et al., 1992; Bhatia et al., 2013), excluding locations with fewer than five individuals to ensure statistical robustness. Isolation by Distance (IBD) was assessed among samples in the Rhine and its sub-catchments by correlating genetic distance (*F*_ST_/1-*F*_ST_) with geographic distance (shortest geographic distance along the Swiss river network (https://www.swisstopo.admin.ch/de/landschaftsmodell-swisstlmregio, Federal Office of Topography swisstopo)). Significance was tested using a Mantel test. Pairwise kinship coefficients were estimated using the KING algorithm as implemented in PLINK 2. To provide a measure of genetic dissimilarity between pairs of individuals among all species, distances were also generated based on allele counts with PLINK 1.9. Before phylogenetic maximum likelihood inference, the LD-pruned data was filtered to retain only SNPs with at least one homozygous individual for the reference and the alternate alleles and was then converted to a PHYLIP file using vcf2phylip v.2 (Ortiz, 2019). Phylogenetic relationships were inferred using RAxML-NG 1.2.2 (Kozlov et al., 2019) using the GTR+ASC_LEWIS substitution model.

Genetic diversity was evaluated with nucleotide diversity (π), observed and expected heterozygosity (*H*o, *H*e) using vcftools. Nucleotide diversity was estimated in 10 kb windows for locations with five or more individuals. Heterozygosity estimates were normalised by dividing heterozygote counts by the total genome size. Inbreeding was assessed by estimating the inbreeding coefficients (*F*_IS_) with vcftools for locations with five or more individuals and runs of homozygosity (ROH). Individual ROH were calculated using the full dataset with the hidden Markov model approach in bcftools. Allele frequencies were calculated from each species and only called genotypes were used for ROH estimation. Individual inbreeding coefficients (FRoH) were calculated by dividing the total length of ROH >300 kb by the total genome size. Finally, contemporary effective population size (*N*e) was estimated with currentNe (Santiago et al., 2024). CurrentNe uses chromosome numbers to correct for non-independence among SNPs and estimates the number of full siblings in the dataset, improving the accuracy of *N*e estimates in species with complex mating systems. The full dataset was downsampled to include just under 2 million SNPs, which is the limit of currentNe. *N*e was estimated only for localities with at least 10 individuals. The genetic indicators (π, *H*o, *F*_ST_, FRoH, *F*_IS_, *N*e) were tested for correlation with the size of the sampled waterbody using Spearman’s or Pearson’s correlation coefficient, depending on normality. The average values of the genetic indicators and kinship across localities were compared between *A. anatina* and *A. cygnea* using Wilcoxon rank-sum tests. Unless otherwise stated, all data manipulation and visualisation of results were performed in R v4.4.2 (Wickham et al., 2019) using Tidyverse packages (R Core Team, 2024) and Rstudio 2024.09.1+394’ (Posit team, 2024). ChatGPT-4o was used to assist in debugging R and Bash scripts, and to refine scientific English after initial drafting by the authors.

## Results

### High-quality draft genome of *A cygnea* and fragmented draft genome of *A. anatina*

Two PacBio libraries from the same *A. cygnea* individual were constructed and sequenced on two SMRT cells, yielding a total of ∼50.4 Gb of HiFi data. The HiFi reads had a mean length of 15,593 bp and 15,916 bp for the two libraries, indicating optimal read quality and fragment length. GenomeScope analysis estimated a genome-wide heterozygosity level of 0.34% (Table 2). The assembled draft genome was of high quality, with a total size of ∼2.46 Gb distributed across 2,008 scaffolds, with a N50 of 2.7 Mb (Table 2). BUSCO completeness scores for Mollusca were relatively low (C: 83.5%), however the completeness score for the broader Metazoa database was substantially higher (C: 96%), similar to other high-quality published Unionid genomes (Table 2).

**Table 2:**
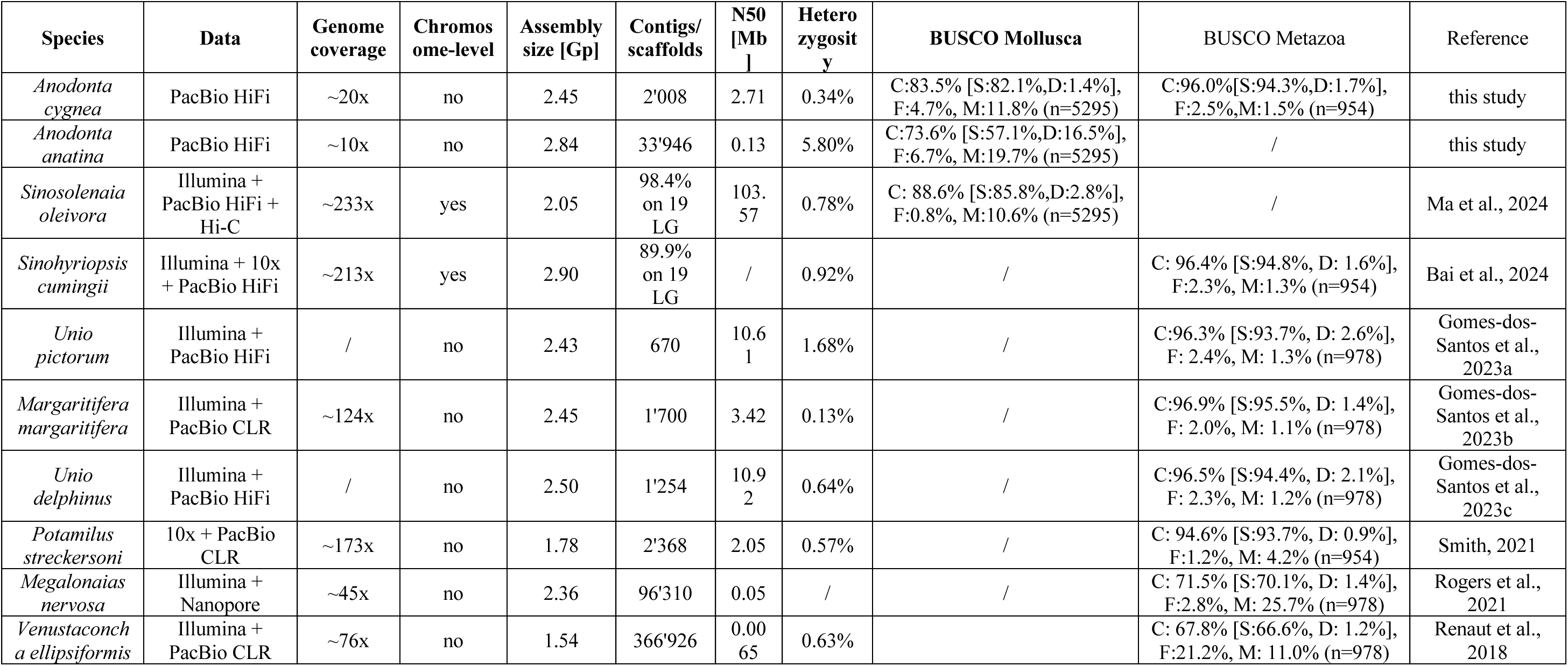
Statistics of the *Anodonta cygnea* and *Anodonta anatina* draft genomes generated in this study and comparison with published genomes of freshwater mussels (order Unionida).

Despite numerous attempts, only two PacBio libraries of *A. anatina* could be successfully constructed and sequenced on a single SMRT cell, resulting in a limited amount of sequencing data available for this species (a total of ∼26.5 Gb of HiFi data). With a mean length of 7,019 bp and 6,954 bp for both libraries, HiFi reads were relatively short. Given the low genome coverage (∼10.6x) and the short read length, the assembly statistics were not of high quality (Table 2). Consequently, the GenomeScope model fit was poor, producing a very low genome size estimate (552-566 Mb) and a very high heterozygosity (4.1-7.4%). The primary HiFi assembly was highly fragmented, consisting of nearly 34,000 contigs with a total length of ∼2.8 Gb, and BUSCO scores were low (C: 73.6% for Mollusca) (Table 2). Nevertheless, due to the relatively high genetic divergence between *A. anatina* and *A. cygnea* (Lopes-Lima et al., 2021, 2024) and the challenges associated with using a distant reference genome for population parameter inference (Thorburn et al., 2023; Maurstad et al., 2024), we opted to use the *A. anatina* draft genome for estimating genetic EBVs for *A. anatina*. While the reference genome is fragmented, this is expected to have only a minor impact on genetic EBVs estimates, with the possible exception of FRoH (see below).

### Whole-genome sequencing uncovers an undescribed *Anodonta* species South of the Alps

Following sampling, we successfully sequenced 421 *Anodonta* specimens collected from 31 locations across Switzerland and assigned them to species based on COI barcodes: 253 *A. anatina*, 151 *A. cygnea*, and 17 *A. exulcerata* (Table 1; Fig. 1A). Specimens were collected from a diverse range of habitats, including large and small lakes, ponds, a river, and a stream, highlighting the broad ecological niches of these species. *A. anatina* was the most abundant species, but *A. cygnea* frequently co-occurred with it at several localities (Fig. 1A). *Anodonta exulcerata* was restricted to Ticino, the region of Switzerland located south of the Alps. Phylogenetic analysis revealed a deep divergence between *A. anatina* and a clade comprising the closely related species *A. cygnea* and *A. exulcerata* (Fig. 1D). This pattern was further supported by PCA, in which over 60% of the total genetic variation separated these two groups (Fig. 1B). Within the *A. anatina* group, we identified a highly divergent population from Locarno (Ticino), that was genetically distinct from all other *A. anatina* individuals (Fig. 2A). This population formed a separate cluster in the PCA (Fig. 1C, 2B). In the nuclear phylogenetic tree, the Locarno population appeared as a monophyletic lineage, exhibiting a level of genetic divergence from other *A. anatina* populations slightly lower than between *A. cygnea* and *A. exulcerata* (Fig. 1D). Further phylogenetic analysis of the COI gene, incorporating *A. anatina* sequences from the literature (Table S3)(Froufe et al., 2017), showed that eight COI sequences clustered with the Italian+Ebro clade, while two sequences clustered with the European clade (Fig. S1). Given this mitochondrial signal and the substantial nuclear divergence, we propose that the Locarno population (Italian+Ebro clade) represents an undescribed *Anodonta* species. Therefore, we refer to this population as *Anodonta* sp. throughout the remainder of this manuscript. Interestingly, two *Anodonta* sp. individuals carried an *A. anatina* haplotype, one of these individuals having about 25% of its genome assigned to *A. anatina* (Fig. 2D). Taken together, this suggests recent hybridization between *Anodonta* sp. and *A. anatina*.

**Figure 1:**
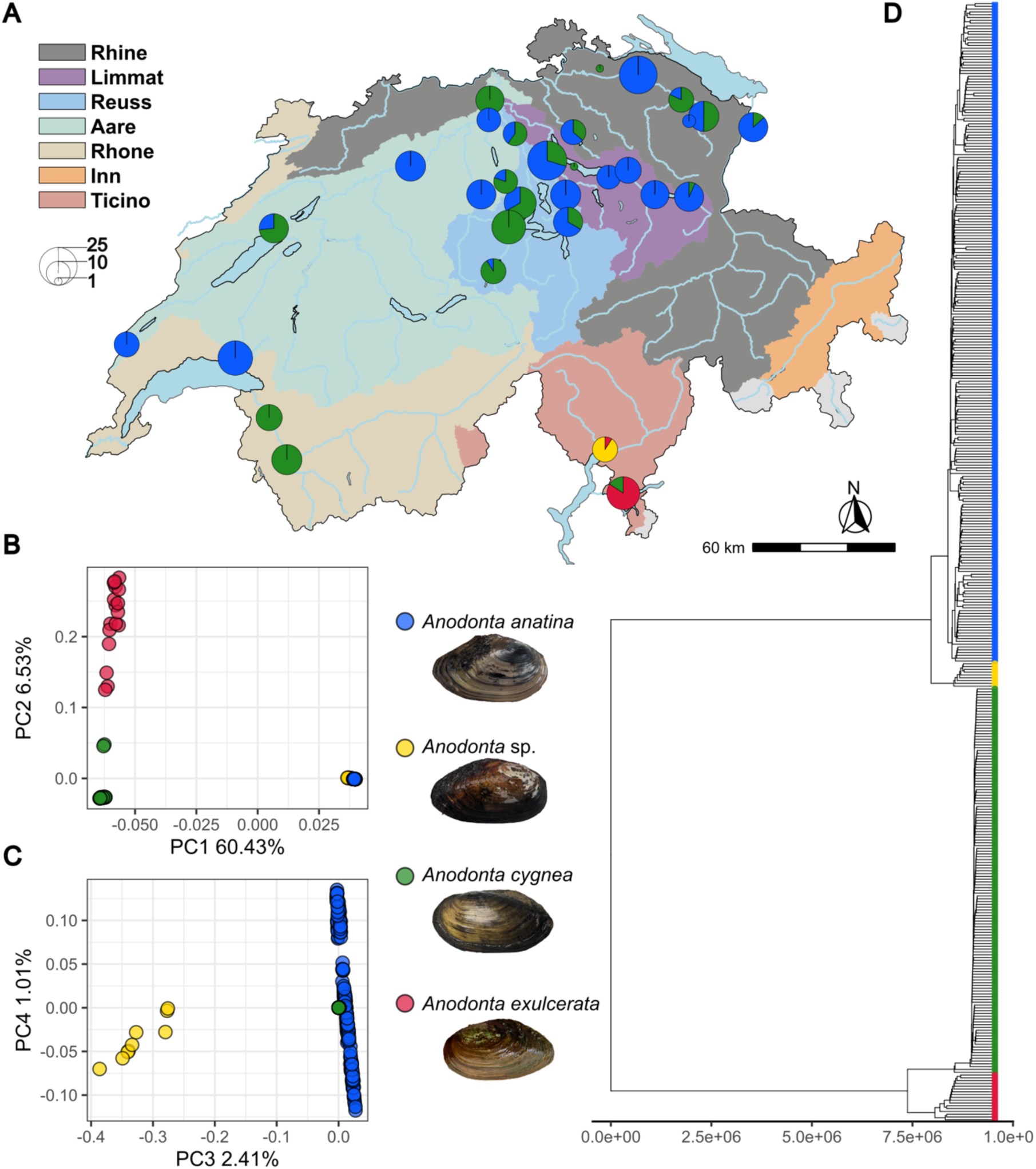
Geographic distribution and genomic divergence of *Anodonta* freshwater mussels in Switzerland. **(A)** Sampling sites and the number of individuals collected in this study. **(B)** Genomic principal components analysis (PCA), showing the first two principal components (PCs) across the four *Anodonta* species. PC1 distinguishes *A. cygnea* and *A. exulcerata* from *A. anatina* and *Anodonta* sp. **(C)** Third and fourth principal components of the genomic PCA, with PC3 separating the undescribed species *Anodonta* sp. **(D)** Genomic divergence among the four species, inferred from allele counts, providing a measure of genetic dissimilarity between pairs of individuals.

**Figure 2:**
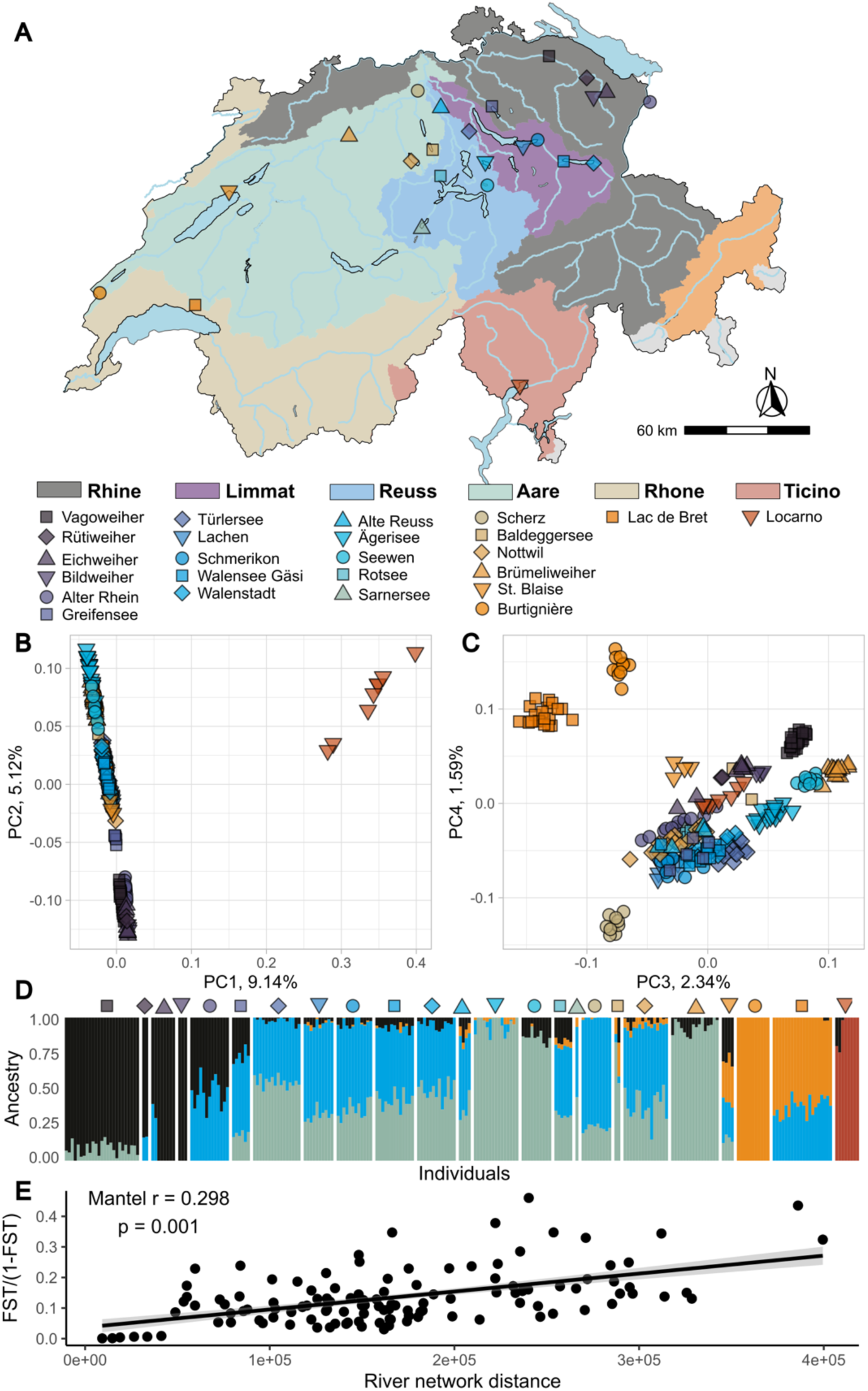
Population structure of *Anodonta anatina and Anodonta sp.* in Switzerland. **(A)** Sampling sites of *A. anatina* within their respective river catchments. **(B)** Genomic principal components analysis (PCA), showing the first two principal components (PCs) across *A. anatina* and *Anodonta* sp. populations. PC1 differentiates *A. anatina* from *Anodonta* sp. **(C)** Third and fourth principal components of the genomic PCA, further distinguishing *A. anatina* populations. **(D)** Clustering analysis using ADMIXTURE, identifying five genetic clusters. Population codes correspond to panel A. **(E)** Significant isolation-by-distance among *A. anatina* populations.

### Population structure follows river catchment in *A. anatina*

Admixture analysis of *A. anatina* identified five well-supported genetic clusters, including the *Anodonta* sp. population from Locarno (Fig. 2D, S2). These groups largely corresponded to major catchment areas: Ticino, Rhône, and Rhine, while populations from the Limmat, Reuss, and Aare catchments were more closely related, reflecting their geographic proximity (Fig. S3). Notably, most populations were clustered by sampling locality, indicating significant genetic structure across populations (Table S4). Most of the populations were genetically distinct, with pairwise *F*_ST_ values ranging from 0.0008 to 0.31, excluding comparisons with *Anodonta* sp. from Ticino (which ranged from 0.22 to 0.35) (Table S4; Fig. S4-S6). Two populations of the Aare catchment were particularly divergent (Fig. 2C): (1) Burtignière, the only river population sampled in this study, which was also geographically distant from other populations in the Aare catchment, and (2) Scherz, a very small stream population. Due to their small size and relative isolation, both populations likely experienced increased genetic drift, leading to their divergence. Furthermore, we detected significant isolation by distance, with geographically distant populations also showing greater genomic differentiation (Fig. 2E). Finally, kinship estimates among populations were very low, indicating that most of the individuals among localities were not related (Table S5).

### Genetic diversity is correlated with waterbody size in *A. anatina*

We next investigated genetic diversity (heterozygosity and nucleotide diversity), inbreeding, and effective population sizes across all *A. anatina* populations (Fig. 3). Observed heterozygosity varied substantially among populations, mean *H*o ranging from 0.0012 to 0.0018 (Fig. 3A). Similarly, nucleotide diversity, calculated as the genome-wide average in 10 kb windows, ranged from 0.0015 to 0.0022 (Fig. 3C). Notably, both measures of genetic diversity were positively correlated with waterbody size, indicating that populations in large lakes exhibited higher genetic diversity compared to those in small ponds (Fig. 3B, D). The inbreeding coefficient (*F*_IS_)—which measures the deficit or excess of heterozygotes relative to Hardy-Weinberg expectations—was generally close to zero, suggesting no strong deviations from Hardy-Weinberg equilibrium (Fig. 3E, F). The exception was the *Anodonta* sp. population from Locarno (Ticino), which exhibited a negative *F*_IS_, indicating an excess of heterozygotes. However, given that only one population of this species was analyzed, and that the use of a heterospecific reference genome can introduce a bias toward excess heterozygosity (Maurstad et al., 2024), this result should be interpreted with caution and is not explored further here. To assess individual inbreeding more directly, we estimated FRoH, the proportion of the genome in long runs of homozygosity (ROH) (Fig. 3G, 3H, S7). While some variation was observed among populations, individual inbreeding appeared to be minimal, with only small ROH and FRoH values below 0.018. However, these estimates should be interpreted cautiously, as the fragmented nature of the *A. anatina* draft genome likely limits the detection of very long ROH. Finally, we estimated effective population size (*N*e), restricting calculations to locations with at least 10 sampled individuals to minimize bias. Estimates were uniformly low, ranging from 5 to 57 (Fig. 3I). Contrary to genetic diversity, *N*e was not significantly correlated with waterbody size, although this may reflect the limited number of populations analyzed (Fig. 3J). Finally, average kinship estimates among individuals within each population were mostly low, ranging from 0.03 to 0.27 (Fig. S8).

**Figure 3:**
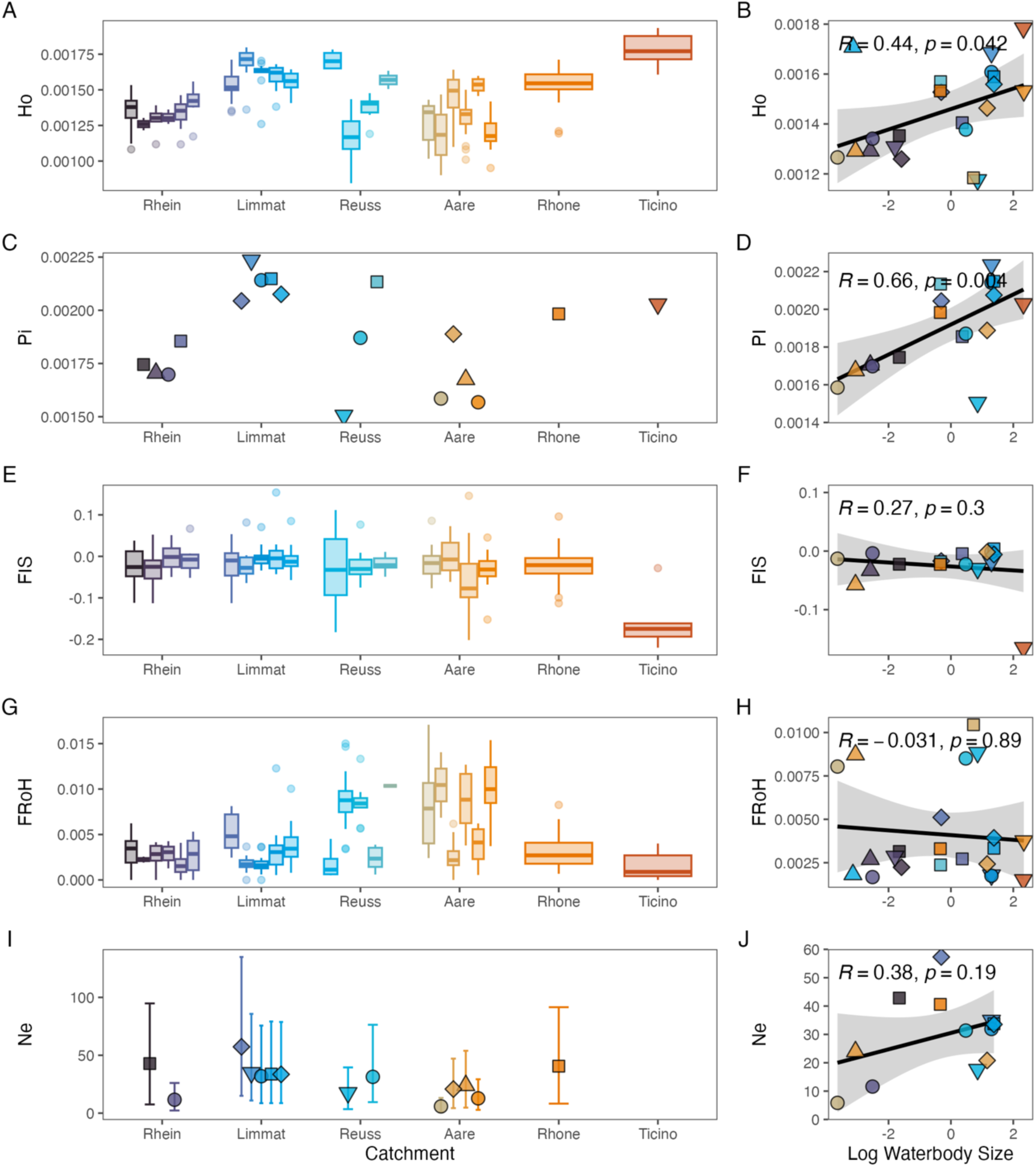
Genetic essential biodiversity variables in *Anodonta anatina and Anodonta sp*. **(A)** Individual observed heterozygosity per sampling locality (all localities included). **(B)** Observed heterozygosity is positively correlated with waterbody size. **(C)** Genome-wide nucleotide diversity (mean over 10 kb windows) per sampling locality, including only sites with at least five individuals. **(D)** Nucleotide diversity is positively correlated with waterbody size. **(E)** Inbreeding coefficient (*F*IS) per sampling locality, including only sites with at least five individuals. **(F)** *F*IS is not correlated with waterbody size. **(G)** Individual inbreeding, quantified as the fraction of runs of homozygosity (FRoH), per sampling locality (all localities included). **(H)** FRoH is not correlated with waterbody size. **(I)** Effective population size (*N*e) per sampling locality, including only sites with at least ten individuals. **(J)** *N*e is not positively correlated with waterbody size.

### Ongoing hybridization between *A. cygnea* and *A. exulcerata*

We identified intriguing patterns in the PCA, where some individuals were positioned between the *A. cygnea* and *A. exulcerata* clusters (Fig. 4B; S9). This suggests ongoing hybridization in Agno (Ticino), the only locality of our dataset where both species occur in sympatry (Fig. 1A). Although only three *A. cygnea* individuals were found at this site, their genetic patterns were notable. One *A. cygnea* individual clustered with *A. cygnea* populations from north of the Alps, whereas the remaining two *A. cygnea* individuals, along with three *A. exulcerata* individuals— each assigned to species by COI barcoding—appeared admixed (Fig. 4A, B). Further evidence of admixture was detected in the ancestry analysis, where most *A. exulcerata* individuals displayed a genomic background partially shared with *A. cygnea* (Fig. 4D; S10). Notably, the two admixed *A. cygnea* individuals displayed particularly high levels of observed heterozygosity compared to the rest of the *A. cygnea* population (Fig. 4C). Additionally, we detected elevated nucleotide diversity in *A. exulcerata* (Fig. 4E), exceeding not only that of *A. cygnea* but also surpassing diversity levels found in *A. anatina* (Fig. 3C).

**Figure 4:**
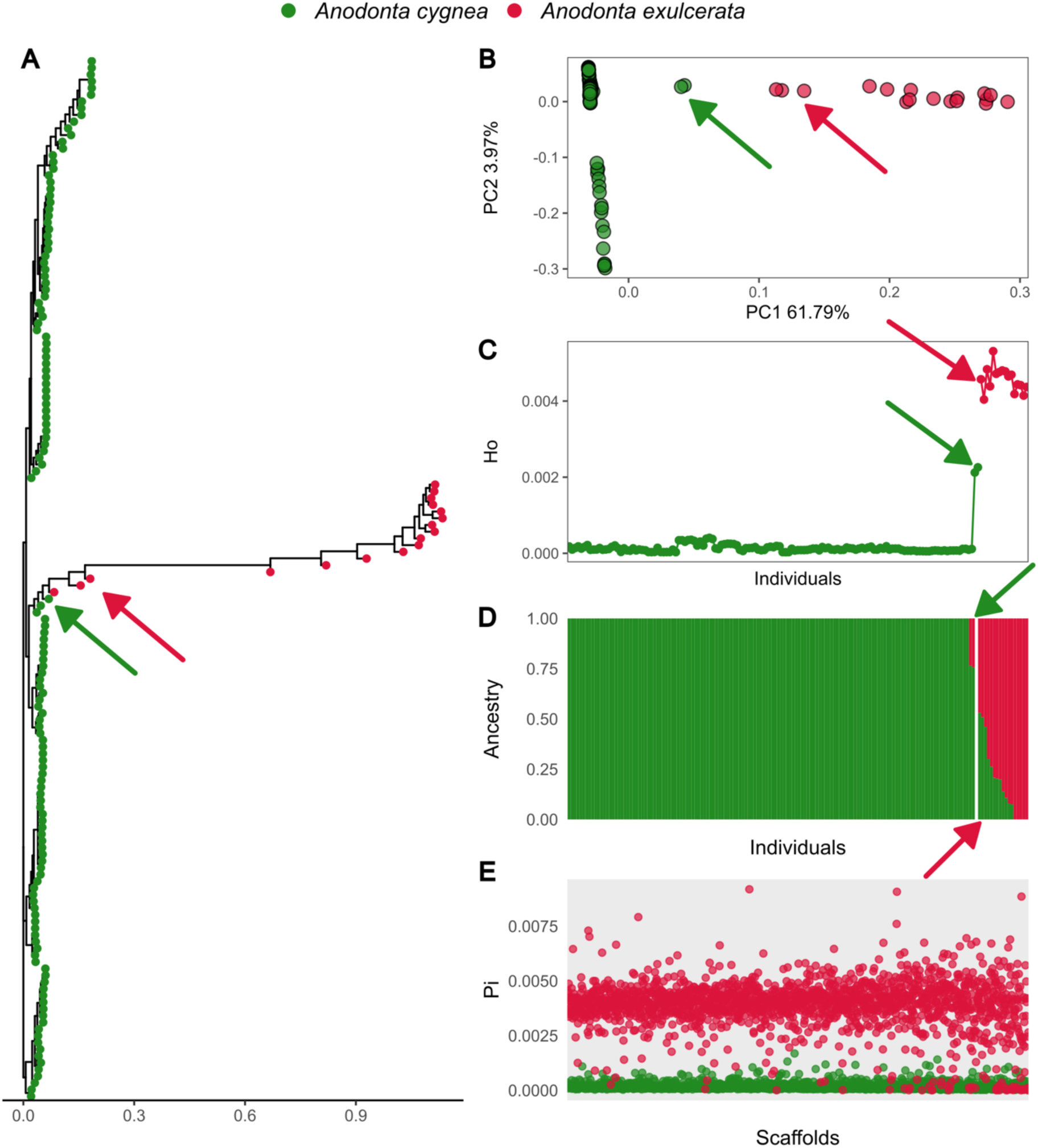
Hybridization between *Anodonta cygnea* and *Anodonta exulcerata*. **(A)** Maximum-likelihood phylogenetic reconstruction highlights hybrid individuals ‘bridging’ *A. cygnea* and *A. exulcerata* (indicated by arrows, same individuals in further panels). **(B)** Genomic principal components analysis (PCA) reveals intermediate individuals between *A. cygnea* and *A. exulcerata*along PC1. **(C)** Individual observed heterozygosity across all *A. cygnea* and *A. exulcerata* individuals. Hybrid *A. cygnea* individuals exhibit intermediate heterozygosity levels between the two species. **(D)** ADMIXTURE analysis shows a significant proportion of the *A. cygnea* genome in most *A. exulcerata* individuals. **(E)** Genome-wide nucleotide diversity (mean over 10 kb windows) per scaffold for each species. Nucleotide diversity differs markedly between species.

### Genomic data suggest facultative selfing in *A. cygnea*

We next examined the population structure of *A. cygnea* in greater detail (Fig. 5A), excluding the two hybrids (Fig. 5B, C). Genetic structure was highly pronounced, as the admixture analysis identified seven well-supported genetic clusters (Fig. 5D; S11). Additionally, two populations (Alte Reuss and Rotsee) were particularly divergent, together accounting for about 22% of the genetic variation in the first axis of the genomic PCA (Fig. 5B). Further principal component axes also differentiated populations from one another (Fig. 5C; S12). The strong genetic structuring was especially evident in the phylogenetic tree, where all individuals clustered within their respective populations (Fig. S13), and in the high genetic differentiation among populations, with *F*_ST_ values ranging from 0.04 to 0.68 (Table S6; Fig. S14). An intriguing finding emerged from the kinship estimates within populations. Under this framework, an individual’s kinship coefficient with itself is 0.5, while full siblings have a coefficient of 0.25. In many populations, the average kinship coefficient exceeded 0.1, indicating that individuals were highly related (Fig. S15, Table S7). Additionally, some individuals from different localities were nearly genetically identical, indicating a reproductive mode other than strict outcrossing (Fig. S13). Furthermore, genetic diversity levels, both in terms of heterozygosity and nucleotide diversity, were particularly low (Fig. 5E, F). While the inbreeding coefficients were not high (Fig. 5G), we observed exceptionally high levels of individual inbreeding, with FRoH values ranging from 0.3 to 0.59 (Fig. 5H). Taken together, these results suggest that *A. cygnea* likely reproduces via facultative selfing. The term “facultative” is appropriate because we observed hybrids with *A. exulcerata*, indicating that *A. cygnea* retains the capacity for outcrossing. Additionally, some genetic variation persisted within populations, which would not be expected in an obligate selfer (Fig. S13). In line with these observations, effective population size (*N*e) estimates were very low, ranging from 4 to 24 (Fig. 5I). Finally, none of the genetic indicators were correlated with waterbody size, except for *F*_IS_ (Fig. S16), and there was no evidence for isolation by distance (Fig. S17).

**Figure 5:**
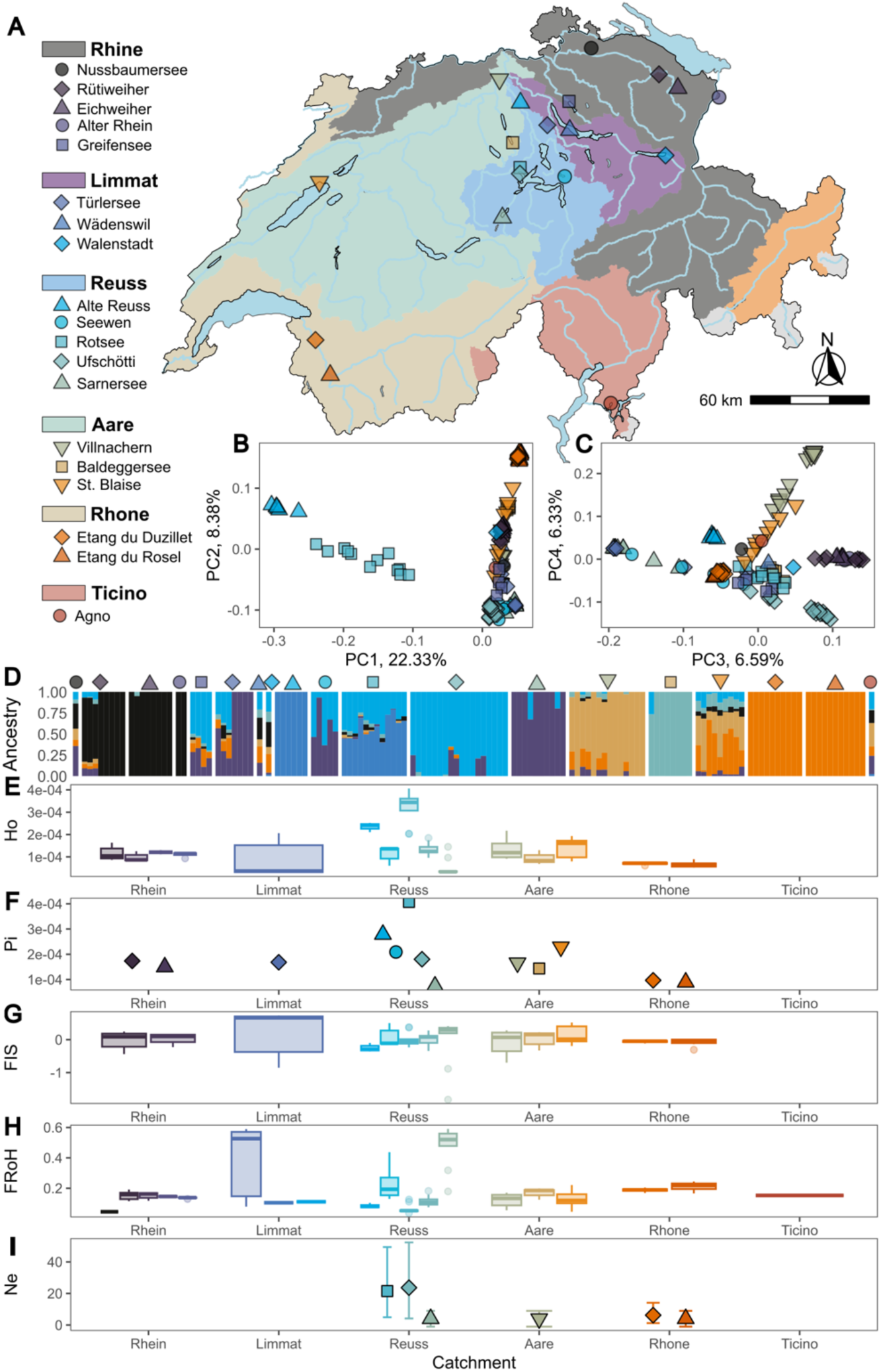
Genetic essential biodiversity variables in *Anodonta cygnea*. **(A)** Sampling sites of *A. cygnea* within their respective river catchments. **(B)** Genomic principal components analysis (PCA), showing the first two principal components (PCs) across *A. cygnea* populations. **(C)** Third and fourth principal components of the genomic PCA, further distinguishing *A. cygnea* populations. **(D)** Clustering analysis using ADMIXTURE, identifying seven genetic clusters. Population codes correspond to panel A. **(E)** Individual observed heterozygosity per sampling locality (all localities included). **(F)** Genome-wide nucleotide diversity (mean over 10 kb windows) per sampling locality, including only sites with at least five individuals. **(G)** Inbreeding coefficient (*F*IS) per sampling locality, including only sites with at least five individuals. **(H)** Individual inbreeding, quantified as the fraction of runs of homozygosity (FRoH), per sampling locality (all localities included). **(I)** Effective population size (*N*e) per sampling locality, including only sites with at least ten individuals.

### Reproductive mode significantly influences genetic indicators

Due to the low number of individuals collected for *A. exulcerata* and *Anodonta* sp., further inferences regarding genetic EBVs in these species were not made. Sufficient sample sizes and geographic coverage were however obtained for *A. anatina* and *A. cygnea*, allowing more detailed comparisons. A striking finding was the substantial differences observed in genetic EBV values between the two species, which differ primarily in their reproductive strategies—*A. anatina* being an obligate outcrosser and *A. cygnea* likely a facultative selfer. In terms of genetic diversity, this change in reproductive mode was associated with a 10-fold reduction in nucleotide diversity and 11-fold reduction in observed heterozygosity, both of which were highly significant (Fig. 6A, B). Regarding genetic differentiation, *A. cygnea* exhibited on average three times as high genetic differentiation (*F*_ST_) between locations and four times as high relatedness within each location (Fig. 6C, D). In terms of inbreeding, the shift in reproductive mode increased individual inbreeding (FRoH) by a factor of 36, while the population inbreeding coefficients (*F*_IS_) were not significantly differentiated among species (Fig. 6E, F). Finally, effective population size was reduced by approximately threefold in *A. cygnea* (Fig. 6G).

**Figure 6:**
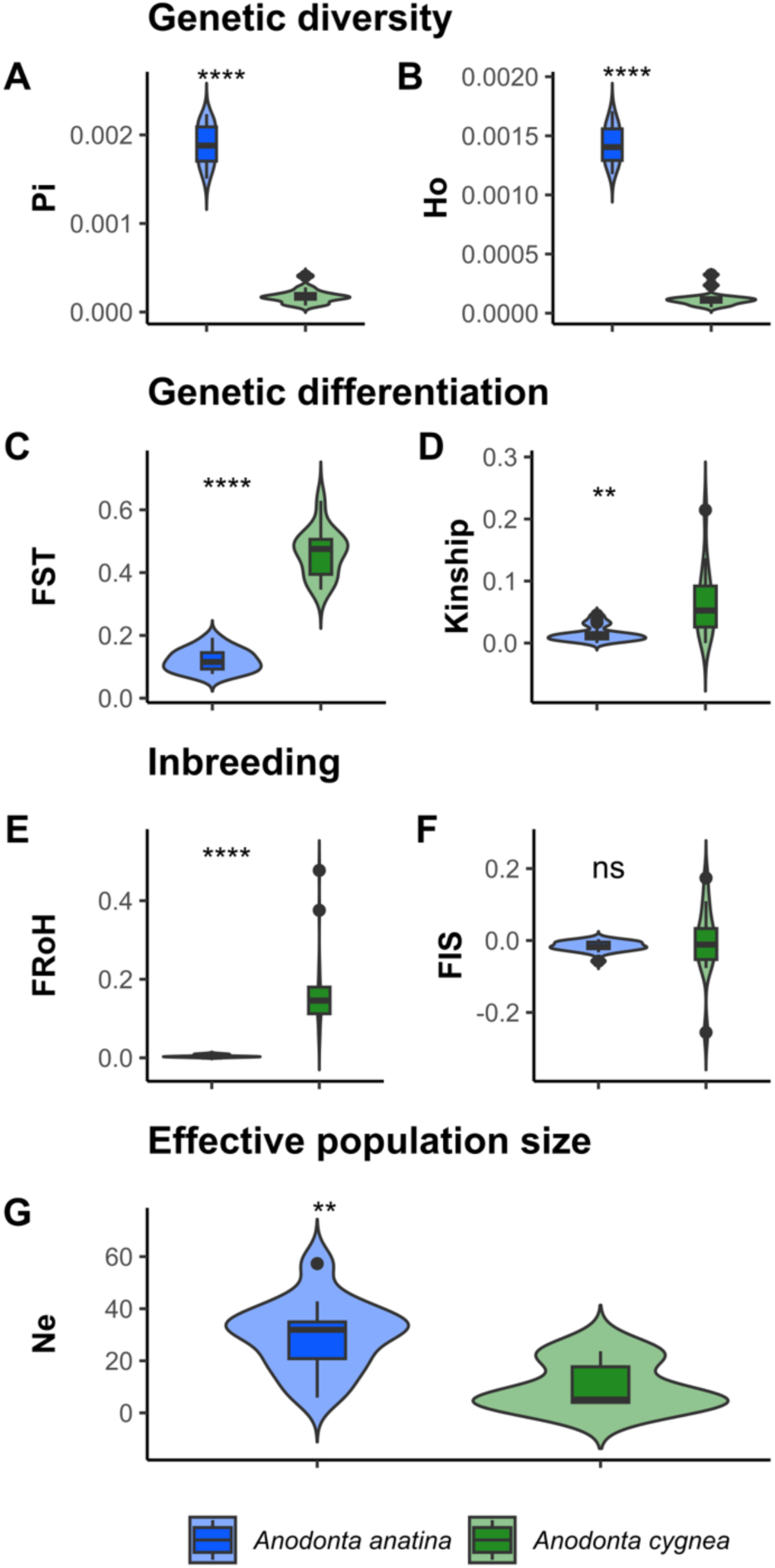
Comparison of genetic indicators between *Anodonta anatina* and *Anodonta cygnea*. **(A)** Nucleotide diversity is significantly higher in *A. anatina* (calculated for sites with at least five individuals; Wilcoxon rank-sum test: *p* < 0.0001). **(B)** Observed heterozygosity is significantly higher in *A. anatina* (calculated for all sites with n>1; Wilcoxon rank-sum test: *p* < 0.0001). **(C)** Genetic differentiation among populations (*F*ST) is significantly lower in *A. anatina* (calculated for sites with at least five individuals; Wilcoxon rank-sum test: *p* < 0.0001). **(D)** The average kinship coefficient within populations is significantly lower in *A. anatina* (calculated for all sites with n>1; Wilcoxon rank-sum test: *p* < 0.01). **(E)** The ROH-based inbreeding coefficient (FRoH) is significantly lower in *A. anatina* (calculated for all sites; Wilcoxon rank-sum test: *p* < 0.0001). **(F)** The inbreeding coefficient (*F*IS) does not differ between *A. anatina* and *A. cygnea* (calculated for sites with at least five individuals; Wilcoxon rank-sum test: *p* > 0.05). **(G)** Effective population size (*N*e) is significantly larger in *A. anatina* (calculated for sites with at least ten individuals; Wilcoxon rank-sum test: *p* < 0.01).

## Discussion

Freshwater mussels play a crucial role in aquatic ecosystems but are among the most endangered aquatic invertebrates. Despite their ecological importance, limited genomic data hinder conservation efforts. In this study, we generated draft genomes for *Anodonta cygnea* and *A. anatina* and resequenced 421 individuals across Switzerland. Our findings reveal key genetic insights, including the discovery of an undescribed *Anodonta* species south of the Alps, strong genetic structuring of *A. anatina* by catchment area, and reduced genetic diversity in small pond populations, indicating higher vulnerability. Notably, we show that a shift in the reproductive mode of *A. cygnea,* most likely facultative selfing, drastically lowers genetic diversity and effective population sizes, increasing extinction risk. These results highlight the need to integrate reproductive strategies into conservation planning and emphasize the importance of habitat connectivity and genetic monitoring for freshwater mussel conservation.

### New genomic resources for European unionids

While the essential role of reference genomes in biodiversity conservation has been well established (Formenti et al., 2022; Theissinger et al., 2023), genomic resources for freshwater mussels remain scarce, with only eight published genomes currently available (Table 2). By providing two additional genome assemblies, our study contributes to filling this gap. Despite considerable efforts, the two draft genomes differ in their quality, mainly caused by the difference in sequencing coverage that was achievable. Interestingly, the assembly quality metrics of the *A. cygnea* genome were comparable to other published unionid genomes, despite lower sequencing coverage (Table 2). This suggests that high-quality assemblies can be obtained with a modest PacBio HiFi coverage, making draft genome sequencing more cost-effective. Furthermore, comparisons with previously published genomes indicate that achieving chromosome-scale assemblies requires additional technologies (e.g. Hi-C) and that solely increasing PacBio coverage has only a marginal impact on assembly contiguity. Finally, our findings suggest that the Mollusca Benchmarking Universal Single-Copy Orthologs (BUSCO) database may not be fully appropriate for Unionids, which likely exhibit true gene losses. Even the chromosome-scale reference genome of *Sinosolenaia oleivora* had a BUSCO completeness score of only 88.6% (Ma et al., 2024), highlighting the need to revise the Mollusca database of single-copy orthologs, as previously suggested for Patellids (Halstead-Nussloch et al., 2024).

### An undescribed *Anodonta* species south of the Alps

Our results reveal cryptic diversity in the Ticino catchment area, south of the Alps, where we identified a population of *Anodonta* sp., representing a currently undescribed species. The potential existence of cryptic species within *A. anatina* has been previously suggested, where studies have documented five divergent mitochondrial lineages, of which at least three had corresponding nuclear divergence, supporting their recognition as distinct evolutionary significant units (ESUs) (Froufe et al., 2014, 2017; Lopes-Lima et al., 2016). Our findings confirm that one of these ESUs, the Ebro & Italy clade, should be considered a separate species based on significant nuclear divergence highlighted here. Interestingly, we also uncovered genomic signatures of hybridization between *A. anatina* and *Anodonta* sp., highlighting that these genomes are still able to recombine, similar to *A. cygnea* and *A. exuclerata*. Although we did not conduct a detailed morphological analysis, it is reasonable to assume that *A. anatina* and *Anodonta* sp. lack diagnostic morphological differences, given their recent evolutionary divergence and the well-documented morphological plasticity of this genus (Riccardi et al., 2020). Finally, as our study is based on a single population, future research should prioritize comprehensive sampling across the full geographic range of this lineage before any formal taxonomic revision can be proposed.

### Genetic essential biodiversity variables for *Anodonta anatina*

A key finding regarding genetic diversity in *A. anatina* is that both nucleotide diversity and heterozygosity were positively correlated with water body size, suggesting that populations inhabiting small ponds are more vulnerable than those in large lakes, due to their lower adaptive potential. A positive correlation between genome-wide heterozygosity and geographic range was also recently found in Felidae (Meeus et al., 2025). Analysis of genetic differentiation revealed strong structuring by catchment area and isolation by distance, with most populations also being significantly differentiated from one another, emphasizing the species’ metapopulation structure. Because mussel dispersal primarily occurs via glochidia parasitizing host fish, our findings highlight the importance of maintaining connectivity among water bodies by avoiding barriers that restrict fish movement. *A. anatina* is considered a host generalist, capable of parasitizing multiple fish species, though studies suggest a preference for *Perca fluviatilis* (Huber and Geist, 2019). Inbreeding levels were generally low, indicating genetic mixing among diverse individuals. However, estimates of the fraction of runs of homozygosity (FRoH) should be interpreted cautiously due to the fragmented nature of the draft genome and the low sequencing coverage (Silva et al., 2024). Our estimates of effective population size (*N*e) were low, ranging from 20 to 60, well below the recommended threshold of *N*e > 500 for maintaining long-term evolutionary potential (Traill et al., 2010; Jamieson and Allendorf, 2012). Most populations fell below 50, the critical short-term threshold to avoid inbreeding depression, suggesting that *A. anatina* populations face a high extinction risk. These low *N*e values may be the result of recent population bottlenecks after the intense period of eutrophication freshwater ecosystems faced in the 1960s-1980s (Vonlanthen et al., 2012). We acknowledge that our *N*e estimates may be underestimated due to the relatively small sample sizes, and that *N*e remains a challenging metric to estimate accurately (Waples, 2025). Yet, even if the *N*e of small connected lakes and ponds within each catchment were combined— assuming that other unsampled populations have similar sizes—it is unlikely that the metapopulation *N*e would reach 500. To complement genetic estimates, direct assessments of census population size (*Nᴄ*) would be valuable. Since all mussels in this study were marked before release, a recapture experiment in the coming years could provide insights into *Nᴄ* and allow comparison with *N*e estimates, as the *N*e/*Nᴄ* ratio is increasingly recognized as an important metric in applied conservation (Waples, 2024).

### Hermaphroditism and selfing: reproductive advantage with evolutionary costs

To date, *Anodonta cygnea* is the only unionid species in Europe described as predominantly hermaphrodite (Lima et al., 2012). *A. exulcerata* may also exhibit hermaphroditism, as it shares with *A. cygnea* the mitochondrial H-ORF, a genetic feature characteristic of hermaphroditic species that have lost the doubly uniparental inheritance system (Breton et al., 2011, 2018; Chase et al., 2018; Riccardi et al., 2020). However, the genetic diversity within the single *A. exulcerata* population was significantly higher than *A. cygnea,* suggesting a lack of self-fertilization. Similarly, while *A. anatina* has been reported as facultative hermaphrodite, self-fertilization has not yet been directly investigated (Hinzmann et al, 2013). The evolution of hermaphroditism has been hypothesized to be favoured in situations where mates are sparsely distributed and access to compatible gametes is limited (Ghiselin, 1969; Eppley and Jesson, 2008). In freshwater mussels, hermaphroditism is thought to be advantageous in low-density populations with limited water flow, where opportunities for gamete dispersal are constrained (Bauer, 1987; Stewart et al., 2013). This reproductive strategy provides an immediate benefit by ensuring reproductive success in the absence of mates (Jarne and Charlesworth, 1993). However, hermaphroditism—particularly when associated with high levels of selfing—carries evolutionary costs, including rapid decline in genetic diversity, and consequently, reduced adaptive potential. Evolutionary theory predicts that obligate self-fertilizing hermaphrodites represent evolutionary dead ends, which was supported by meta-analyses (Stebbins, 1957; Takebayashi and Morrell, 2001). Consistent with this prediction, obligate hermaphrodites among freshwater mussels appear to have evolved relatively recently, with gonochorism (separate sexes) representing the ancestral state (Stewart et al., 2013). One possible mechanism for mitigating extinction risk is facultative selfing, combined with occasional hybridization with closely related species, as observed here between *A. cygnea* and *A. exulcerata*. However, given that *A. cygnea* has a much broader geographic distribution than *A. exulcerata*, such genetic exchange is likely restricted to a few localized sympatric populations. As a result, across its broader range, *A. cygnea* is likely at a greater long-term risk of extinction than *A. anatina*, despite both species having similar pan-European distributions.

We cannot rule out the possibility that the unusual genomic signal observed in *A. cygnea* results from facultative parthenogenesis, in which embryos develop from unfertilized eggs (Smith and Shuker, 2025). In particular, terminal fusion automixis can drastically reduce heterozygosity within a single generation (Lampert, 2009; Lehtonen et al., 2012), potentially explaining the genomic pattern resembling clonality observed in some *A. cygnea* populations (Fig. S13). Furthermore, the presence of sperm was not specifically investigated in the ‘self-fertilized’ individuals examined (Bloomer, 1940, 1943). However, facultative parthenogenesis is less common in hermaphrodites. To minimize stress on the mussels, we did not record individual sex in this study, but literature suggests that *A. cygnea* is predominantly hermaphroditic (Lima et al., 2012). Further studies specifically designed to differentiate selfing from parthenogenesis are needed to confirm this species’ reproductive mode conclusively.

### Hybridization and atypical reproductive modes challenge genetic EBV assessment

Our study highlights the complexities of using genetic EBVs for conservation, particularly in species that exhibit hybridization and atypical reproductive strategies. We detected ongoing hybridization between *A. cygnea* and *A. exulcerata*, with at least five strongly admixed individuals identified in a sympatric population. Notably, this *A. exulcerata* population exhibited unexpectedly high genetic diversity, exceeding even that of *A. anatina*. As only a single population was sampled, it remains unclear whether this elevated diversity is an inherent feature of the species. However, a study on mitochondrial *COI* across the entire *A. exulcerata* range found that *A. exulcerata* had higher haplotype diversity than *A. cygnea* (Froufe et al., 2017). Hybridization appears to be a common phenomenon in freshwater mussels (Sano et al., 2022; Lopes-Lima et al., 2024). More broadly, hybridization is known to play a significant role in the evolution of biodiversity (Taylor and Larson, 2019): for instance, by facilitating adaptation or driving speciation (Meier et al., 2017), but it can also lead to negative outcomes such as outbreeding depression (Frankham et al., 2011) or genetic swamping (Todesco et al., 2016). These dual effects complicate the interpretation of EBVs, as elevated genetic diversity in hybrid populations does not necessarily equate long-term stability or resilience (Quilodrán et al., 2020).

Beyond hybridization, reproductive mode also plays a crucial role in shaping genetic EBVs. Our study provides an ideal system to investigate this, as *A. anatina* and *A. cygnea* differ primarily in their reproductive strategies while sharing similar dispersal abilities, geographic distribution and evolutionary histories (Lopes-Lima et al., 2017). This parallel allows us to attribute the observed differences in genetic EBVs primarily to their reproductive mode. We found that the change in reproductive mode of *A. cygnea* is associated with markedly reduced genetic diversity, increased population structure and inbreeding, and lower effective population sizes. Further investigation is needed to assess whether *A. cygnea* experiences inbreeding depression or can persist with consistently low genetic diversity, and whether alternative mechanisms such as phenotypic plasticity, epigenetic regulation (Venney et al., 2023; Balard et al., 2024), and/or the purging of deleterious mutations (Arunkumar et al., 2015) are alleviating its genetic constraints. The interplay between selfing and hybridization complicates conservation assessments. While these reproductive strategies have been extensively studied in plants, increasing evidence suggests they are also widespread among animals (Avise, 2015; Sperling and Glover, 2023). Defining appropriate thresholds for genetic indicators in species with atypical reproductive modes remains an open question. The recently launched COST Action *Genetic Nature Observation and Action* (https://www.cost.eu/actions/CA23121/) aims to address this challenge, specifically in the Objective 3.4 of the Working Group 3. Ultimately, our study underscores the critical need to integrate species-specific reproductive biology into conservation assessments to ensure that biodiversity monitoring and management are both accurate and effective.

## Conclusion

Our study demonstrates the power of genomics to uncover cryptic diversity, assess population structure, and inform freshwater mussel conservation. We identified an undescribed *Anodonta* species south of the Alps, highlighting the need for taxonomic and genetic assessments before conservation actions. Strong population structuring in *A. anatina* underscores the importance of maintaining hydrological connectivity, while the low genetic diversity in small pond populations signals increased vulnerability. Whole-genome resequencing provides critical insights into genetic diversity, effective population size, and inbreeding, offering a baseline and early warnings before demographic declines are evident. However, genetic indicators must be interpreted cautiously in species with complex reproductive strategies like self-fertilization and hybridization. Genetic monitoring should extend beyond single time points to track trends and detect early signs of genetic erosion (Schwartz et al., 2007; Hoban et al., 2014; Leroy et al., 2018). Global conservation initiatives for freshwater mussels are currently underway (Aldridge et al., 2023) and include habitat restoration, captive breeding, and reintroduction programs (Ferreira-Rodríguez et al., 2019; Geist et al., 2023; Sousa et al., 2023). Breeding programs, successful in other mussel species (Geist et al., 2021, 2023), could aid restoration but must consider genetic risks such as inbreeding or outbreeding depression. Without genomic resources, translocation efforts risk being ineffective (McLennan et al., 2025). Recent initiatives, such as genetic monitoring frameworks (Hogg et al., 2025), and increased collaboration between researchers and conservation practitioners (Shaw et al., 2024) will help integrate genomics into real-world conservation. Future priorities should include expanding genomic resources, refining genetic EBVs for species with atypical reproductive modes, and strengthening partnerships with conservation stakeholders. Applying genomic insights to management will be critical for ensuring the long-term persistence of freshwater mussel populations and broader biodiversity.

## Acknowledgements

We thank Anna Carlevaro, Pascal Stucki, Jukka Jokela, Christoph Walcher, Morris Galli, Jonathan Weber and Marc Odermatt for their help with mussel sampling. We thank the following cantonal authorities for issuing sampling permits: Canton Aargau: Christian Tesini, Astrid Binder, Corinne Gröli, Bruno Schelbert; Canton Zürich: Nina Gremlich; Canton Bern: Daniel Bernet and Olivier Bessire; Canton Schwyz: Philipp Bünter; Canton Zug: Martin Ziegler; Canton Thurgau: Rolf Niederer; Canton St-Gallen: Michael Kugler; Canton Luzern: Peter Kull and Pro Natura Luzern: Patricia Burri; Canton Obwalden: Josef Hess and Kerstin Maier; Canton Neuchâtel: Robin Berger, Christoph Noël; Canton Vaud: Frédéric Hofman, Nathalie Rausis; Canton Valais: Yann Clavien. We thank Pamela Nicholson and Marion Ernst from the NGSP in Bern for HMW DNA extraction and sequencing of the *Anodonta anatina* genome. Whole-genome resequencing was performed at the FGCZ. Data produced and analyzed in this paper were generated in collaboration with the Genetic Diversity Centre (GDC), ETH Zurich. We are grateful to Niklaus Zemp for bioinformatic support with the *Anodonta cygnea* genome assembly and to Gwyneth Halstead-Nussloch for bioinformatic support with the *Anodonta anatina* genome assembly. We also thank Jukka Jokela and Joana L. Santos for valuable discussions and feedback on this manuscript. This work was funded by an Eawag Discretionary Fund Grant to PGDF and AATW. PacBio HiFi sequencing of *A. cygnea* genome was supported by the Biodiversity Genomics Europe project (Grant no. 101059492) funded by the Horizon Europe programme under the Biodiversity, Circular Economy and Environment call (REA.B.3), and co-funded by the Swiss State Secretariat for Education, Research and Innovation (SERI) under contract numbers 22.00173 and 24.00054 and by the UK Research and Innovation (UKRI) under the Department for Business, Energy and Industrial Strategy’s Horizon Europe Guarantee Scheme.

## Scripts and data availability

The raw Illumina resequencing data of the 421 *Anodonta* individuals have been submitted to the European Nucleotide Archive (ENA) and can be accessed under the BioProject ID PRJEB86155 (accession numbers ERS23745809-ERS23746229). The raw PacBio sequencing data of *A. cygnea* have been submitted to the ENA and can be accessed under the BioProject ID PRJEB77385. The assembled draft genomes of *A. cygnea, A. anatina*, the filtered VCF files, the COI alignment and all scripts used in this study are available on the Eawag database ERIC Open (https://doi.org/10.25678/000DY2) and on github https://github.com/ellikafaust/anodonta. Mussel occurrence data have been reported to the Swiss Center for the Cartography of the Fauna (info fauna) and have been linked with the Global Biodiversity Information Facility (GBIF) and the National database of genetic diversity from populations of wild species (GenDiB - project ID: gendib00000143) available at: https://doi.org/10.15468/p3yn5j.

## Supplementary Tables captions

**Table S1:** Metadata including individual codes and ENA accession numbers of the 421 *Anodonta* individuals collected in this study.

**Table S2:** Filtering parameters and number of variants at each filtering step for the three VCF files used in this study.

**Table S3:** *Anodonta anatina* mitochondrial lineage and GenBank accession numbers of the COI sequences retrieved from Froufe et al, 2017, used for the phylogenetic tree displayed in Figure S1.

**Table S4:** Pairwise *F*_ST_ values (Hudson estimator) among *A. anatina* populations having five individuals or more.

**Table S5:** Pairwise individual kinship estimates among all *A. anatina* and *Anodonta* sp. individuals.

**Table S6:** Pairwise *F*_ST_ values (Hudson estimator) among *A. cygnea* and *A. exulcerata* populations having five individuals per species or more.

**Table S7:** Pairwise individual kinship estimates among all *A. cygnea* and *A. exulcerata* individuals.

