## Supplementary methods and figures for "Beyond genetic indicators: how reproductive mode and hybridization challenge freshwater mussel conservation"

### Supplementary Material

**Table of contents:**

**Supplementary methods:** Draft genomes of *Anodonta cygnea* and *Anodonta anatina* (p. 2-4)
**Figure S1:** Neighbour-Joining phylogenetic tree of the *COI* marker, including 10 *Anodonta* sp. from Locarno (Ticino) and all *COI* haplotypes (AA1–AA18) from the *A. anatina* European (EUR) and Italian (ITA) clades (Froufe et al., 2017) (p. 5)
**Figure S2:** ADMIXTURE cross-validation analysis for *A. anatina* and *Anodonta* sp. together with admixture results for K=1 to K=12, separated by sampling locality. (p. 6)
**Figure S3:** *A. anatina* and *Anodonta* sp. ADMIXTURE results for K=1 to K=12, separated by catchment area. (p. 7)
**Figure S4:** Pairwise $F_{ST}$ values among all *A. anatina* and *Anodonta* sp. populations. (p. 8)
**Figure S5:** Maximum-Likelihood phylogenetic tree of *A. anatina* and *Anodonta* sp. populations, coloured by sampling locality. (p. 9)
**Figure S6:** Maximum-Likelihood phylogenetic tree of *A. anatina* and *Anodonta* sp. populations, coloured by catchment area. (p. 10)
**Figure S7:** Runs of homozygosity in *A. anatina* and Anodonta sp. (p. 10)
**Figure S8:** Population pairwise average kinship estimates among *A. anatina* and Anodonta sp. populations.
(p. 11)
**Figure S9:** Genomic PCA (PC1-PC8) for *A. cygnea* and *A. exulcerata.* (p. 12)
**Figure S10:** ADMIXTURE cross-validation analysis for *A. cygnea* and *A. exuclerata* together with admixture results for K=1 to K=6, separated by sampling locality. (p. 13)
**Figure S11:** ADMIXTURE cross-validation analysis for *A. cygnea* (excluding hybrids) together with admixture results for K=1 to K=10, separated by sampling locality. (p. 14)
**Figure S12:** Genomic PCA (PC1-PC8) for *A. cygnea* (excluding hybrids). (p. 15)
**Figure S13:** Maximum-Likelihood phylogenetic tree of *A. cygnea*, coloured by sampling locality. (p. 16)
**Figure S14:** Pairwise $F_{ST}$ values among all *A. cygnea* and *A. exulcerata* populations. (p. 17)
**Figure S15:** Population pairwise average kinship estimates among *A. cygnea* populations.
(p. 18)
**Figure S16:** Absence of strong association between genetic indicators (observed heterozygosity, inbreeding coefficient, nucleotide diversity, fraction of runs of homozygosity, effective population size) and waterbody size in *A. cygnea.* (p. 19)
**Figure S17:** Absence of isolation by distance in *A. cygnea.* (p. 20)

**Supplementary methods: Draft genomes of *Anodonta cygnea* and *Anodonta anatina***

*Sampling and molecular species identification*

Collection permits were requested in advance to the relevant cantonal authorities of nature protection (Canton Aargau for mussels collected in Scherz; Canton Zürich for mussels collected in Türlersee). Five adult *Anodonta anatina* individuals were hand-collected from a small stream in Scherz, Switzerland (47°26′ 40.20″N 8°11′05.89″E) on the 24$^{th}$ of February 2023 and brought back alive to the Eawag laboratory in Dübendorf, Switzerland. Two adult individuals of the swan mussel *Anodonta cygnea* were collected by snorkeling in Türlersee, Switzerland (47°15'57.2"N 8°30'35.6"E) on the 7$^{th}$ of July 2023 and brought back alive to the laboratory in Dübendorf. All mussels were kept alive in the Eawag experimental facility Aquatikum until processing. They were maintained in an open-water circuit at 19°C and fed twice a week using a mixture of lab-grown algae. Given the morphological plasticity of *Anodonta* mussels, species identification was initially based on shell morphology and subsequently confirmed by sequencing a COI barcode. Briefly, a small tissue biopsy was taken from the foot of each of the seven individual mussels and DNA was extracted using the E.Z.N.A Tissue DNA kit (Omega Bio-tek) according to the manufacturer's instructions. DNA integrity, quantity, and purity were assessed using NanoDrop Eight (thermo scientific; Witec AG). A fragment of the COI mitochondrial gene was amplified with forward primer LCO1490-JJ (5'-3' primer sequence: CHACWAAYCATAAAGATATYGG) and reverse primer HCO2198-JJ (5'-3' primer sequence: AWACTTCVGGRTGVCCAAARAATCA). PCR cycling conditions were: an initial step of 95°C for 15 min, followed by 35 cycles of 94°C for 30 s, 51°C for 90 s, 72°C for 60 s and then a final extension of 72°C for 10 min. Amplified PCR products were purified and Sanger sequenced by Microsynth AG, Balgach, Switzerland. The COI sequences of the seven *Anodonta* individuals were compared to the NCBI GenBank database using the Basic Local Alignment Search Tool (BLAST) to confirm morphological species identification.

*High molecular weight DNA extraction, library preparation and PacBio HiFi sequencing*

For *Anodonta cygnea,* the two individuals were dissected on a glass plate on ice and small tissue pieces were flash frozen in liquid nitrogen. The tissue samples were then sent to the University of Florence, Italy, for High-Molecular Weight (HMW) DNA extraction, library preparation and PacBio HiFi sequencing conducted as part of the European Reference Genome Atlas project Biodiversity Genomics Europe (ERGA-BGE). Tissue leftovers from both individuals were deposited as vouchers at the Natural History Museum in Bern, Switzerland (voucher IDs: NMBE584869 and NMBE584870).

HMW DNA was extracted from the specimen NMBE584870 using the E.Z.N.A. Mollusc & Insect DNA Kit (Omega Bio-Tek) according to the manufacturer's protocol. Nucleic acids yield was quantified in a Qubit fluorimeter using a Qubit dsDNA HS Assay (Life Technologies) and DNA purity was assessed by comparing absorbance values at 260, 280 and 230 nm in a Tecan Infinite M200 Pro spectrophotometer using a NanoQuant plate (Tecan). Integrity of DNA was then assessed by pulsed-field gel electrophoresis. Whole DNA was sheared in a Megaruptor 2 DNA shearing system (Diagenode) using large fragment hydropores with a target mean fragment length of 20 kb. Fragments profile was assessed by capillary electrophoresis on a Fragment Analyzer using an HS large fragment 50 kb kit (Agilent Technologies). Repair and A-tailing of DNA fragments and SMRTbell adapters ligation were performed according to the SMRTbell prep kit 3.0 protocol (Pacific Biosciences). Primer annealing, polymerase binding and preparation of internal DNA control were performed using the Pacific Biosciences Sequel II binding kit and DNA internal control complex 3.2 according to the manufacturer's protocol. Library fragments shorter than 10 kb were removed by eluting libraries in a Blue Pippin automated pulse-field gel electrophoresis device (Sage Science). Sequencing runs were set up using SMRT Link v11.1. Samples were sequenced in HiFi mode in a Pacific Biosciences Sequel IIe platform using Sequel II Sequencing plates 2.0 and two 8M ZMW SMRT cells with a 30-hour movie time and 2 hours of pre-extension time for a target 25X genome coverage.

For *Anodonta anatina*, several HMW DNA extractions were performed in the laboratory at Eawag, Dübendorf. Despite apparent initial good quality and integrity, the DNA was consistently degraded when quality control was performed at the sequencing center (Next Generation Sequencing Platform (NGSP) in Bern, Switzerland), which was likely due to strong DNases remaining in the DNA extractions. Therefore, a specimen (PB107) was brought alive to the NGSP where it was dissected and processed for HMW DNA extraction. Despite multiple tests from different protocols and kits, a satisfactory HMW DNA extraction could not be achieved. Eventually, HMW DNA was extracted using MagAttract HMW DNA Kit (Qiagen) from foot and mantle tissues. There was sufficient DNA to construct two PacBio libraries from foot and mantle tissues, respectively. Prior to SMRTbell library preparation, genomic DNA was assessed for quantity, quality and purity using a Qubit 4.0 flurometer (Qubit dsDNA HS or BR Assay kit; Q32851/Q32850, Thermo Fisher Scientific), an Advanced Analytical FEMTO Pulse instrument (Genomic DNA 165 kb Kit; FP-1002-0275, Agilent) and a Denovix DS-11 UV-Vis spectrophotometer, respectively. SMRTbell libraries were prepared according to the PacBio guideline SMRTbell prep kit 3.0. Briefly, sheared gDNA was concentrated and cleaned using 1 x SMRTbell clean-up beads. The samples were then quantified and qualified to be in the range of 9-20 Kb using a Qubit 4.0 flurometer (Qubit dsDNA HS Assay kit; Q32851, Thermo Fisher Scientific) and an Advanced Analytical FEMTO Pulse instrument (Genomic DNA 165 kb Kit; FP-1002-0275, Agilent), respectively. The rest of the procedure as referenced above was followed including end-repair & A-tailing, ligation of barcoded overhang adapters and then purification of the library using AMPure PB beads as well as a nuclease treatment. The only deviation was that for libraries with ≥1 µg, a gel-based size selection was employed to remove fragments below 10 Kb according to the document entitled: "Technical note GEL CASSETTE SIZE SELECTION METHODS FOR WGS HIFI LIBRARIES", following the protocol for a Sage Science BluePippin device (PacBio part number 102-326-503). Final libraries were checked for quantity and quality using a Qubit 4.0 flurometer (Qubit dsDNA HS Assay kit; Q32851, Thermo Fisher Scientific) and an Advanced Analytical FEMTO Pulse instrument (Genomic DNA 165 kb Kit; FP-1002-0275, Agilent). Instructions in SMRT Link Sample Setup were followed to prepare the SMRTbell library for sequencing (PacBio SMRT Link v12). Shortly, PacBio Sequencing primer v3.2 and Sequel DNA Polymerase 3.0 were annealed and bound, respectively, to the DNA template libraries using a Sequel II Binding Kit 3.2 (PacBio Part number 102-333-300) and the complex was cleaned using SMRTbell clean-up beads. Both libraries were loaded at an on-plate concentration of 120-170pM using adaptive loading, along with the use of Sequel II DNA internal control complex. SMRT sequencing was performed in CCS mode on the Sequel IIe with Sequel Sequencing kit 3.0, SMRT Cells 8M, a 2h pre-extension followed by a 30 h movie time and via PacBio SMRT Link v12. Thereafter, the CCS generation is performed on the Sequel IIe and the read segmentation workflow was run in SMRT Link v12. All steps were performed at the Next Generation Sequencing Platform, University of Bern, Switzerland.

*Genome assembly*

To assess *A. cygnea* genome features, a 21 bp k-mer database was generated with HiFi data (QV≥20) from both cells using the count function of meryl v1.3 (Rhie et al., 2020). K-mer frequencies and genome statistics were obtained using GenomeScope v2.0 (Ranallo-Benavidez et al., 2020). Genome assembly was then conducted using hifiasm v0.19.9 (Cheng et al., 2021), using default parameters and the purging parameter -l set to 1, given the low level of estimated genome-wide heterozygosity (0.34%). Assembly metrics were retrieved using asmstats and assembly completeness was assessed with BUSCO v5.2.2 (Simão et al., 2015). The Mollusca OrthoDB database (odb10; 2024-01-08; n:5295) and the metazoa OrthoDB database (odb10; 2024-01-08; n:954) were used for the analyses.

For the *A. anatina* draft genome, a 31 bp k-mer database was generated from HiFi reads (QV ≥ 20) obtained from two separate libraries (foot and mantle tissue) using the count function of meryl v1.3. The two k-mer databases were merged with the union-sum function of meryl. K-mer frequencies and genome statistics were obtained using GenomeScope v2.0. Genome assembly was then conducted using hifiasm v0.16.1-r375, using default parameters and the purging parameter -l set to 3. Assembly metrics were retrieved using asmstats and assembly completeness was assessed using BUSCO v5.2.2. The mollusca OrthoDB database (odb10; 2020-08-05; n:5295) and the eukaryota OrthoDB database (odb10; 2020-09-10; n:255) were used for the analyses.

**References from supplementary methods:**


Cheng, H., Concepcion, G. T., Feng, X., Zhang, H., and Li, H. (2021). Haplotype-resolved de novo assembly using phased assembly graphs with hifiasm. *Nat. Methods* 18, 170–175.

Ranallo-Benavidez, T. R., Jaron, K. S., and Schatz, M. C. (2020). GenomeScope 2.0 and Smudgeplot for reference-free profiling of polyploid genomes. *Nat. Commun.* 11, 1432. doi: 10.1038/s41467-020-14998-3

Rhie, A., Walenz, B. P., Koren, S., and Phillippy, A. M. (2020). Merqury: reference-free quality, completeness, and phasing assessment for genome assemblies. *Genome Biol.* 21, 245. doi: 10.1186/s13059-020-02134-9

Simão, F. A., Waterhouse, R. M., Ioannidis, P., Kriventseva, E. V., and Zdobnov, E. M. (2015). BUSCO: assessing genome assembly and annotation completeness with single-copy orthologs. *Bioinformatics* 31, 3210–3212.

**Figure S1:** Neighbour-Joining phylogenetic tree of the *COI* marker, including 10 *Anodonta* sp. from Locarno (Ticino) and all *COI* haplotypes (AA1–AA18) from the *A. anatina* European (EUR) and Italian (ITA) clades (Froufe et al., 2017). Based on 100 bootstrap replicates; nodes with ≥90% bootstrap support are shown.

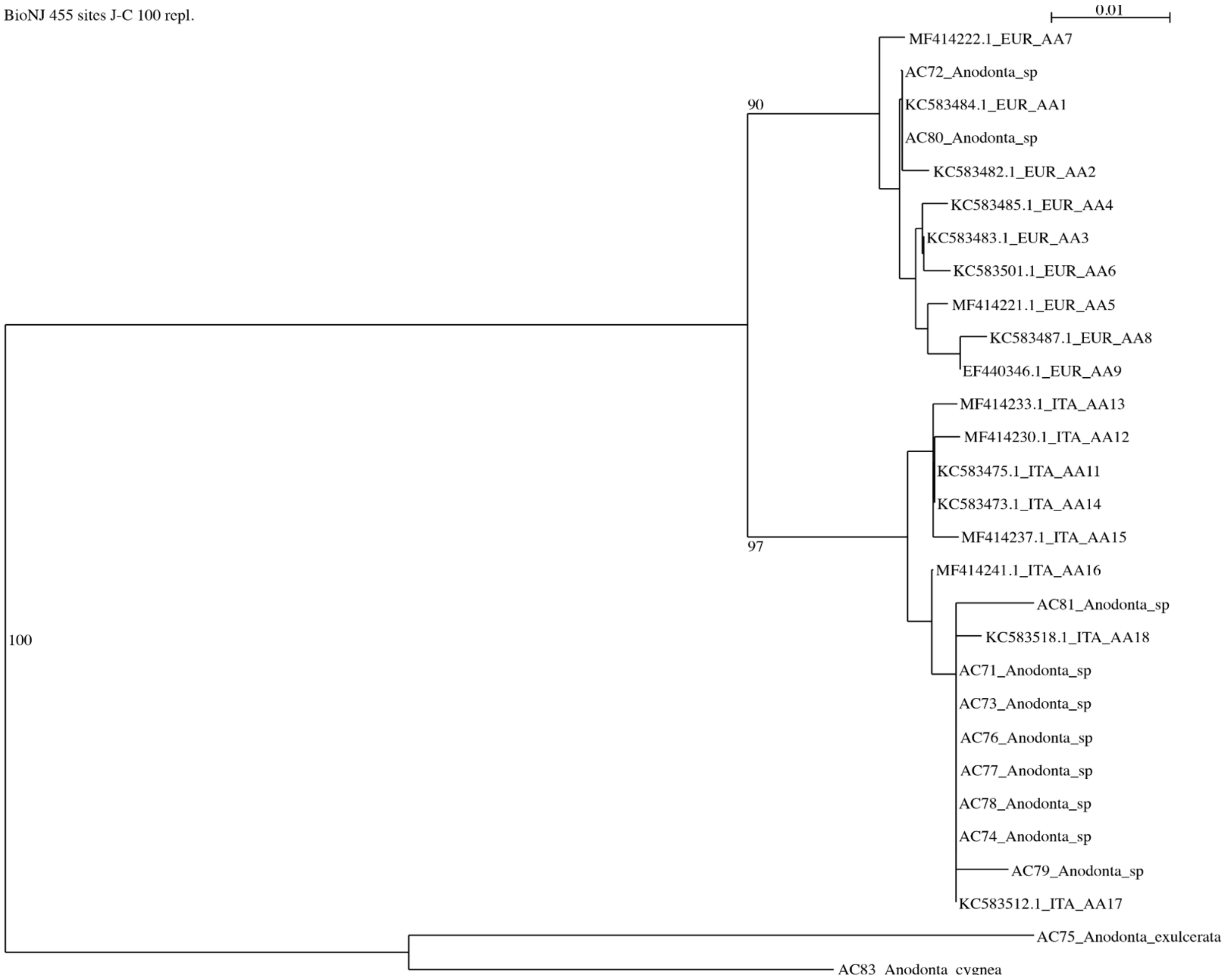

**Figure S2:** ADMIXTURE cross-validation analysis for *A. anatina* and *Anodonta* sp. together with admixture results for K=1 to K=12, separated by sampling locality.

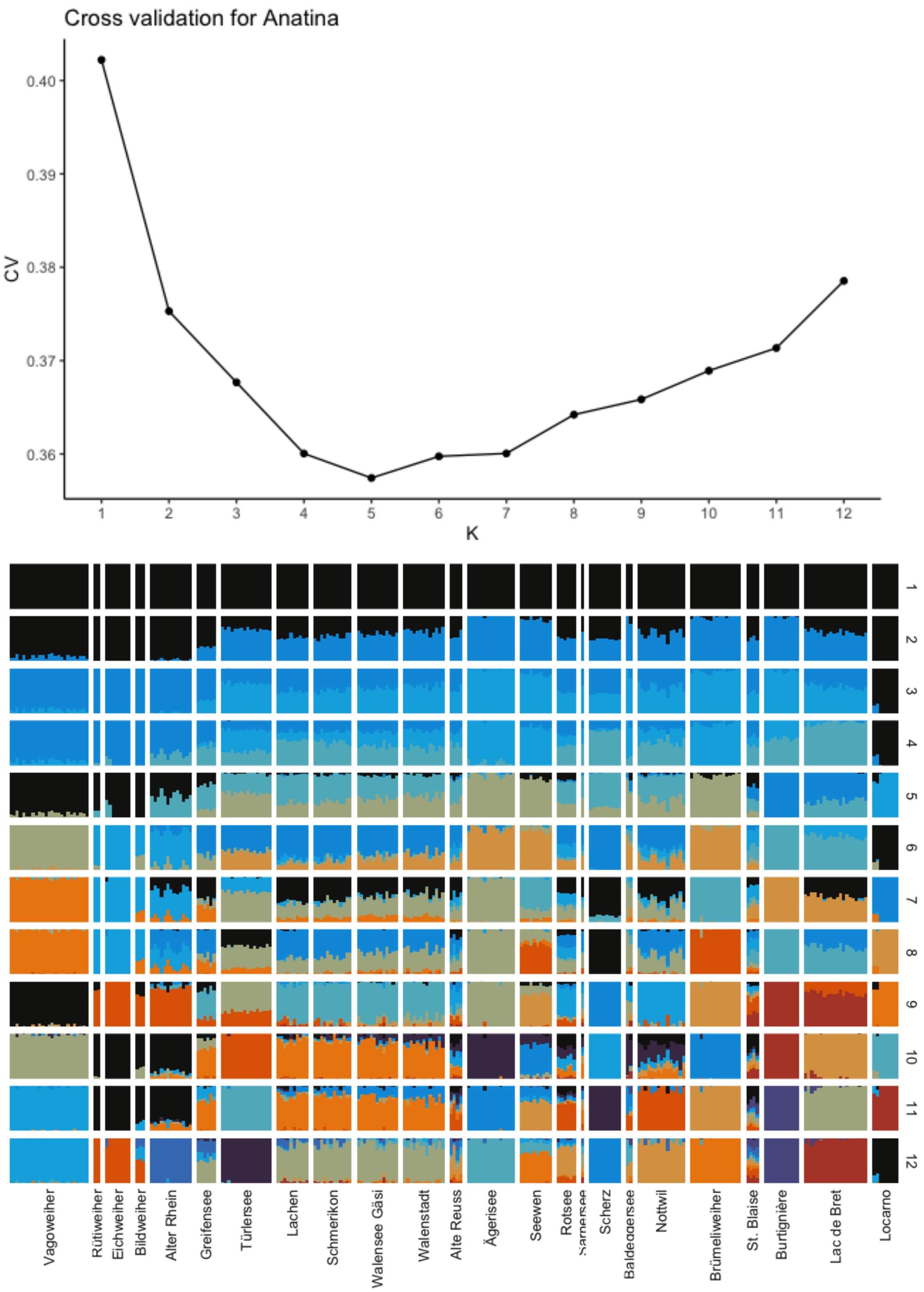

**Figure S3:** *A. anatina* and *Anodonta* sp. ADMIXTURE results for K=1 to K=12, separated by catchment area.

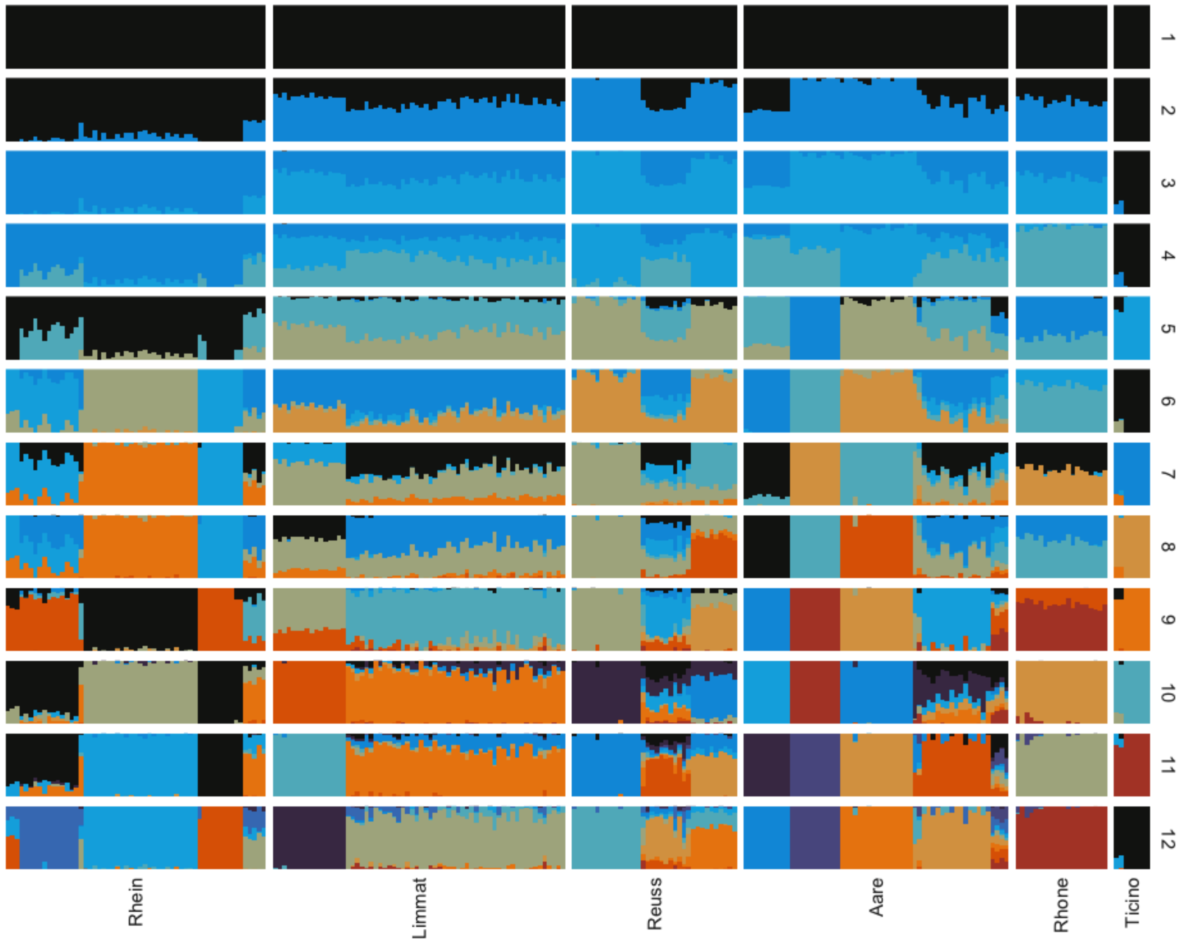

**Figure S4:** Pairwise $F_{ST}$ values among all *A. anatina* and *Anodonta* sp. populations.

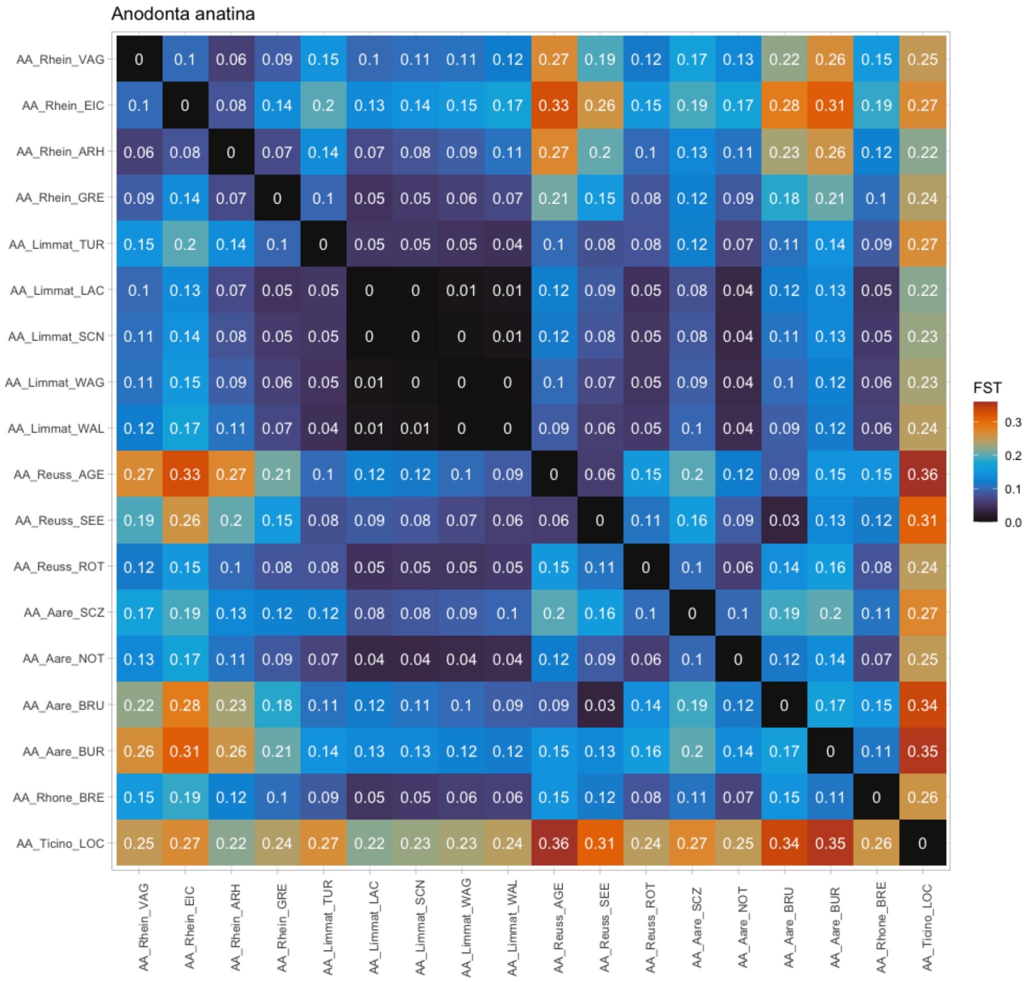

**Figure S5:** Maximum-Likelihood phylogenetic tree of *A. anatina* and *Anodonta* sp. populations, coloured by sampling locality.

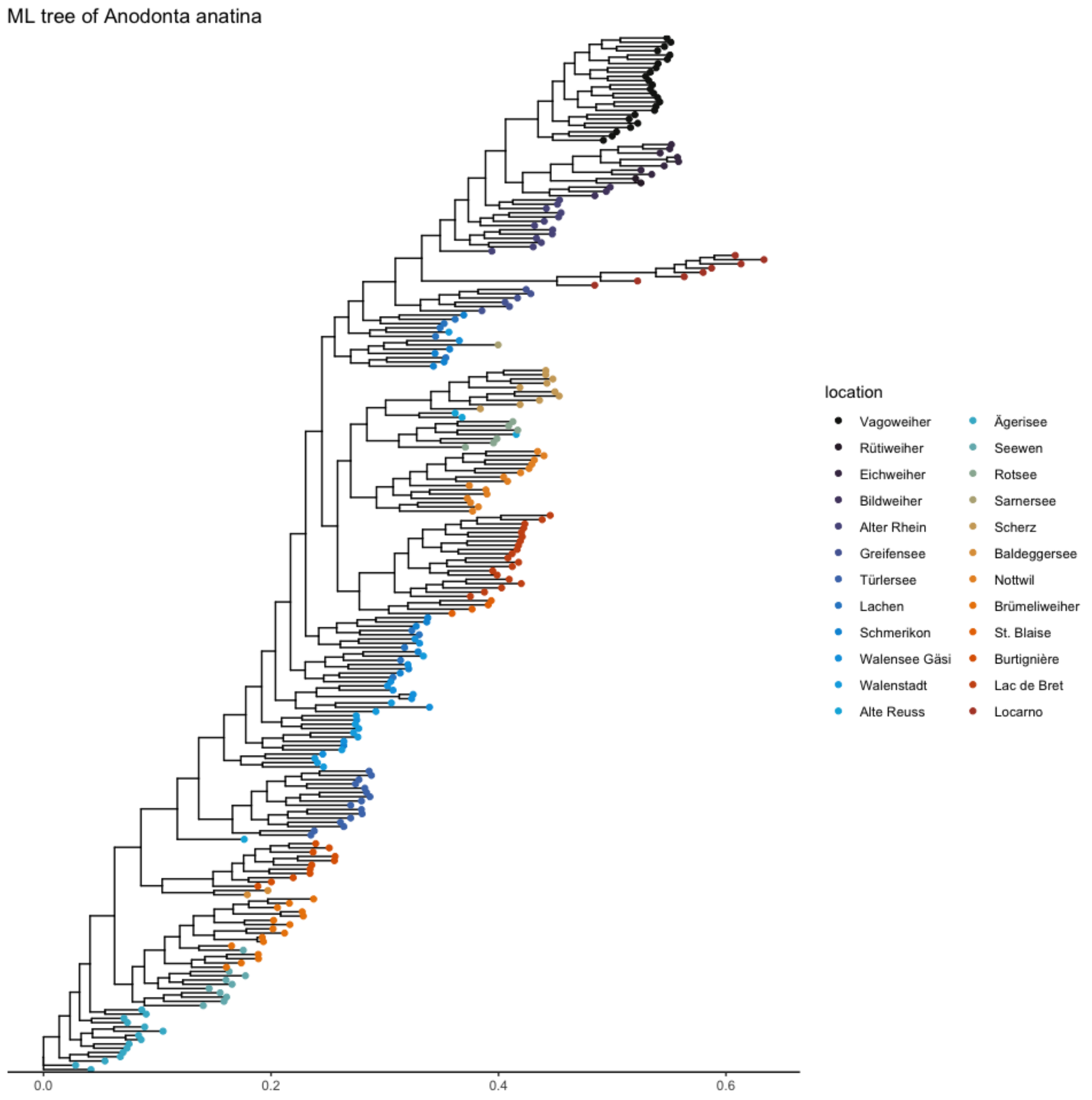

**Figure S6:** Maximum-Likelihood phylogenetic tree of *A. anatina* and *Anodonta* sp. populations, coloured by catchment area.

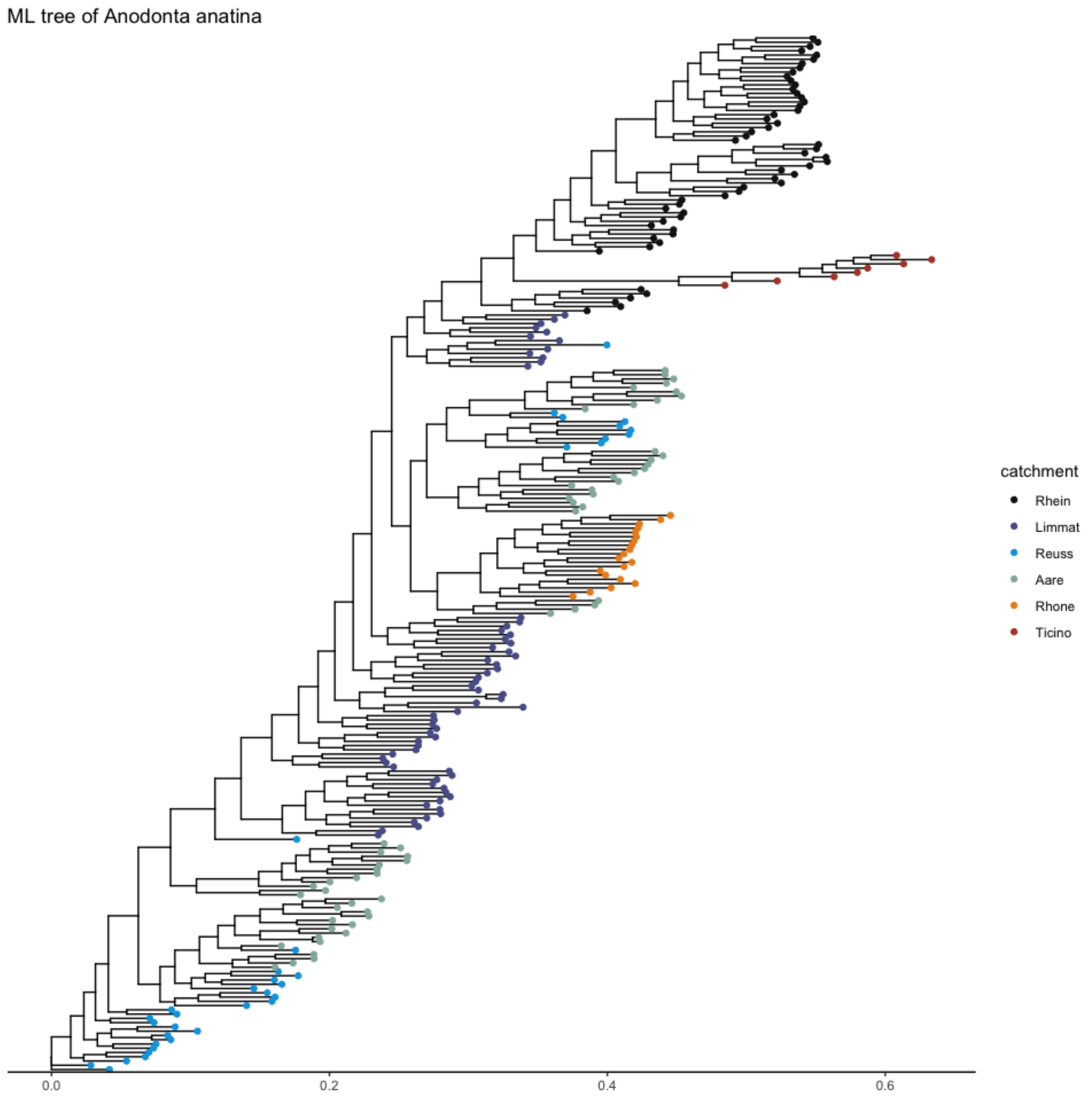


**Figure S7:** Runs of homozygosity in *A. anatina* and *Anodonta* sp.

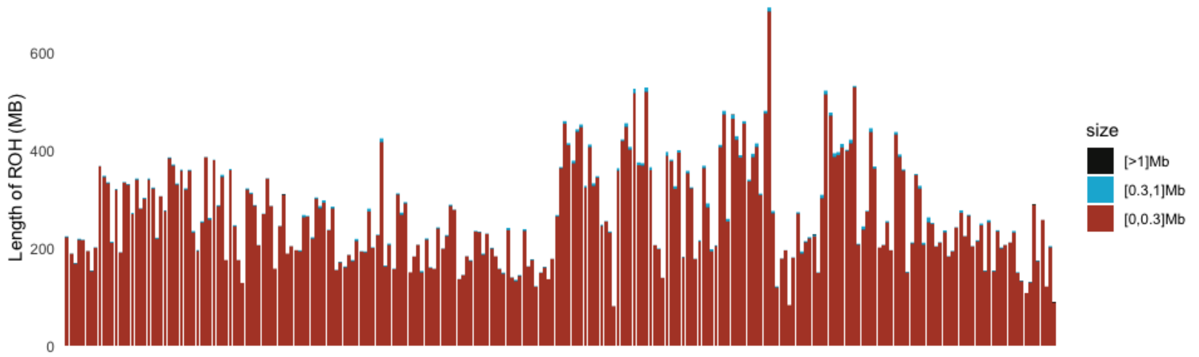

**Figure S8:** Population pairwise average kinship estimates among *A. anatina* and *Anodonta* sp. populations.

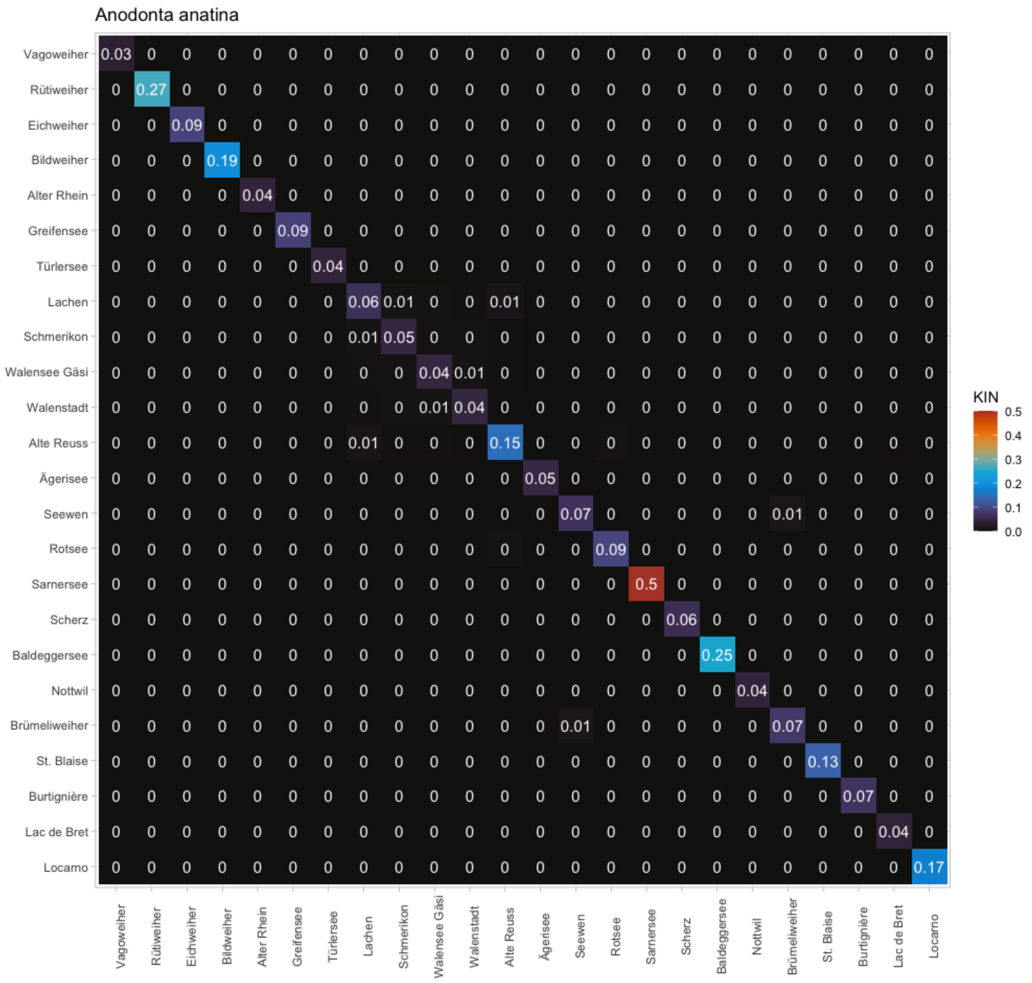

**Figure S9:** Genomic PCA (PC1-PC8) for *A. cygnea* and *A. exulcerata.*

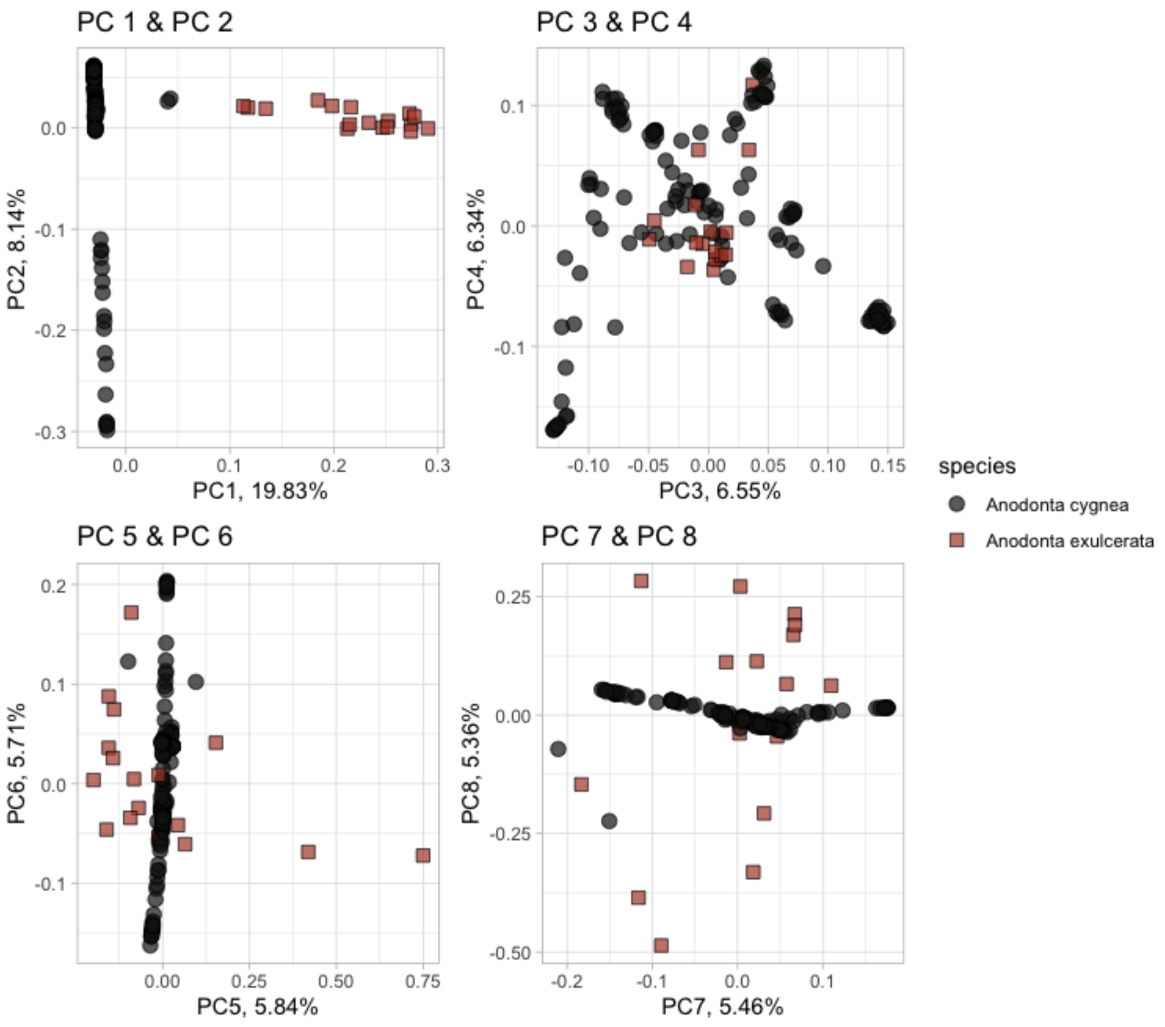

**Figure S10:** ADMIXTURE cross-validation analysis for *A. cygnea* and *A. exuclerata* together with admixture results for K=1 to K=6, separated by sampling locality.

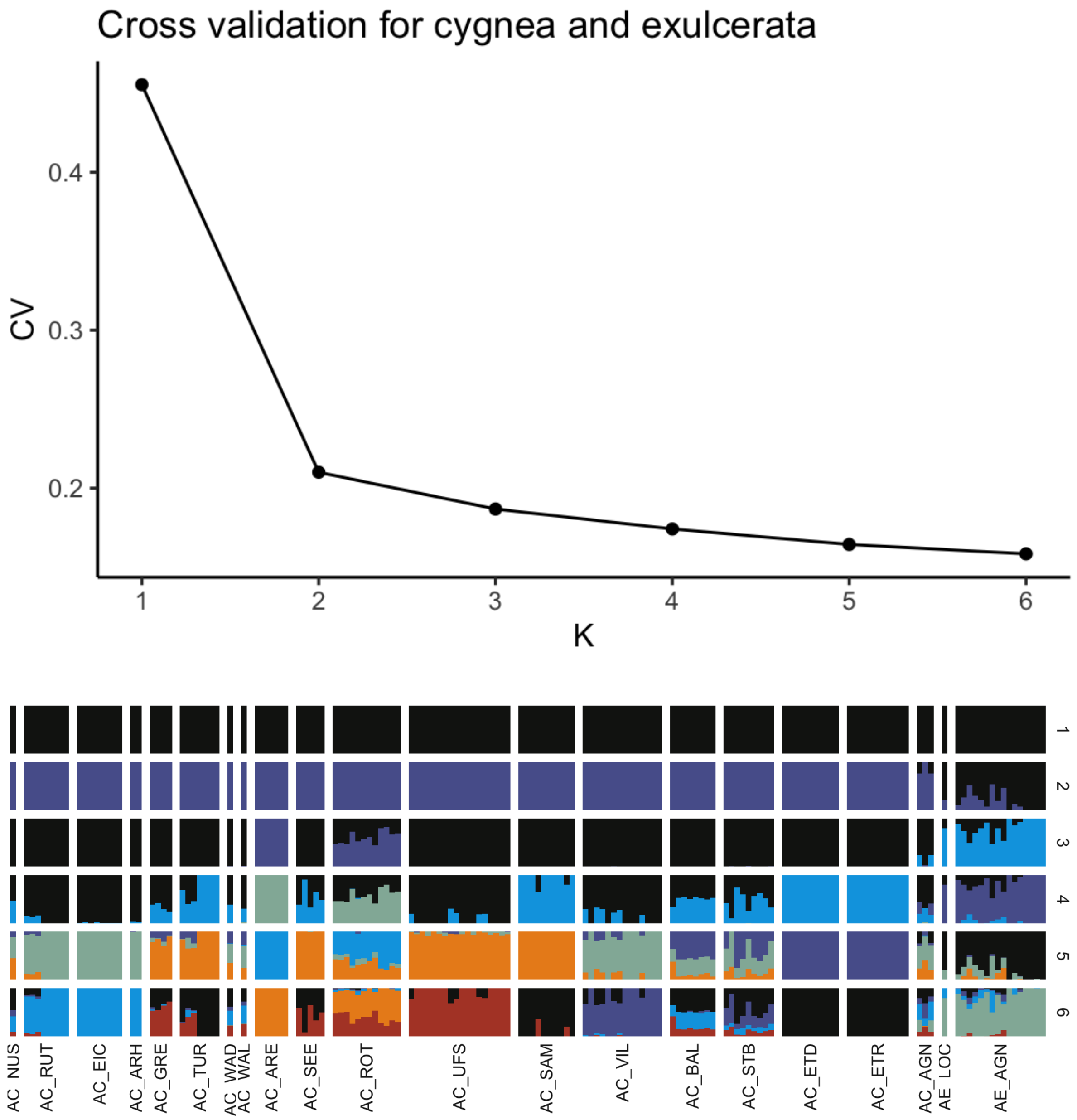

**Figure S11:** ADMIXTURE cross-validation analysis for *A. cygnea* (excluding hybrids) together with admixture results for K=1 to K=10, separated by sampling locality.

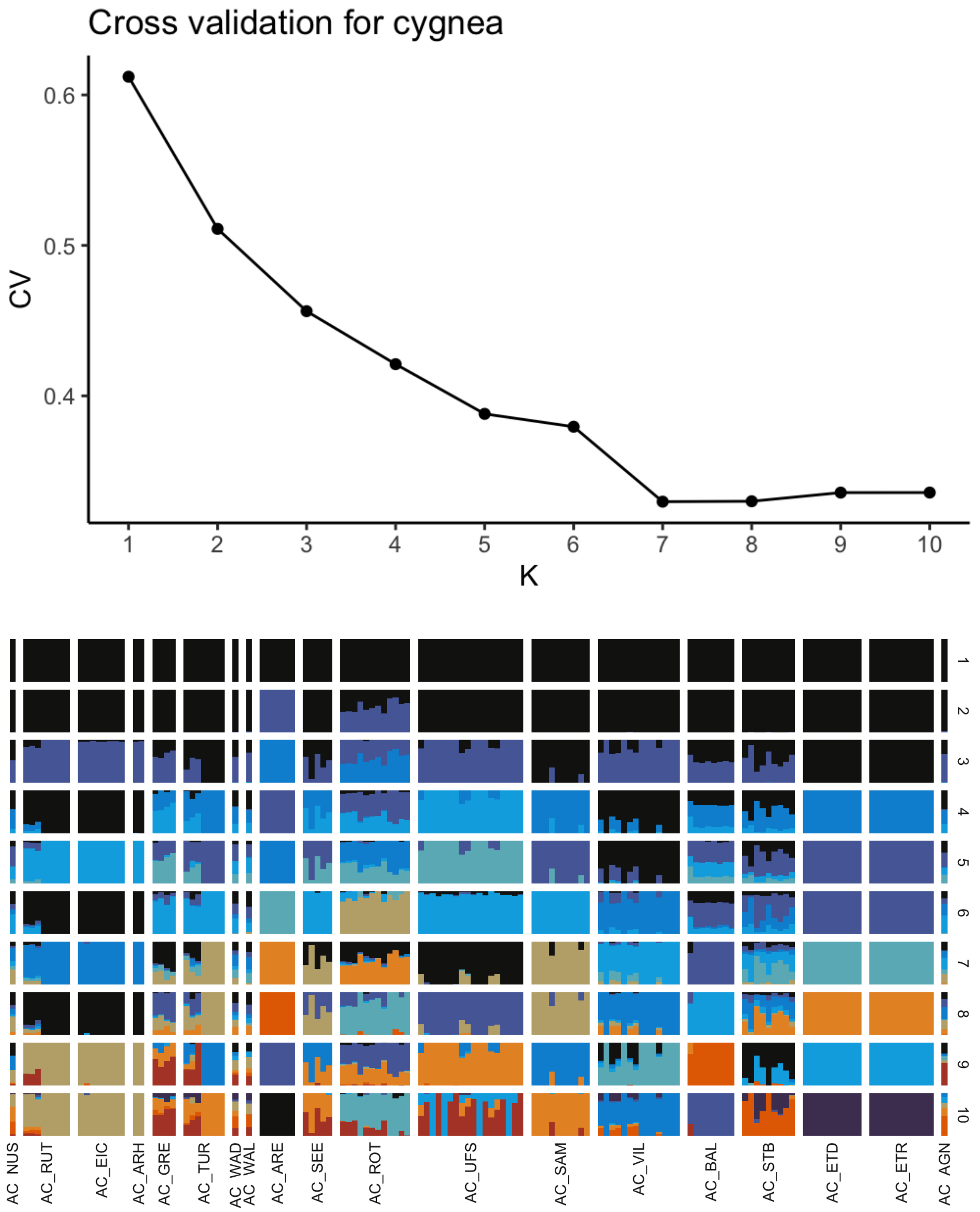

**Figure S12:** Genomic PCA (PC1-PC8) for *A. cygnea* (excluding hybrids).

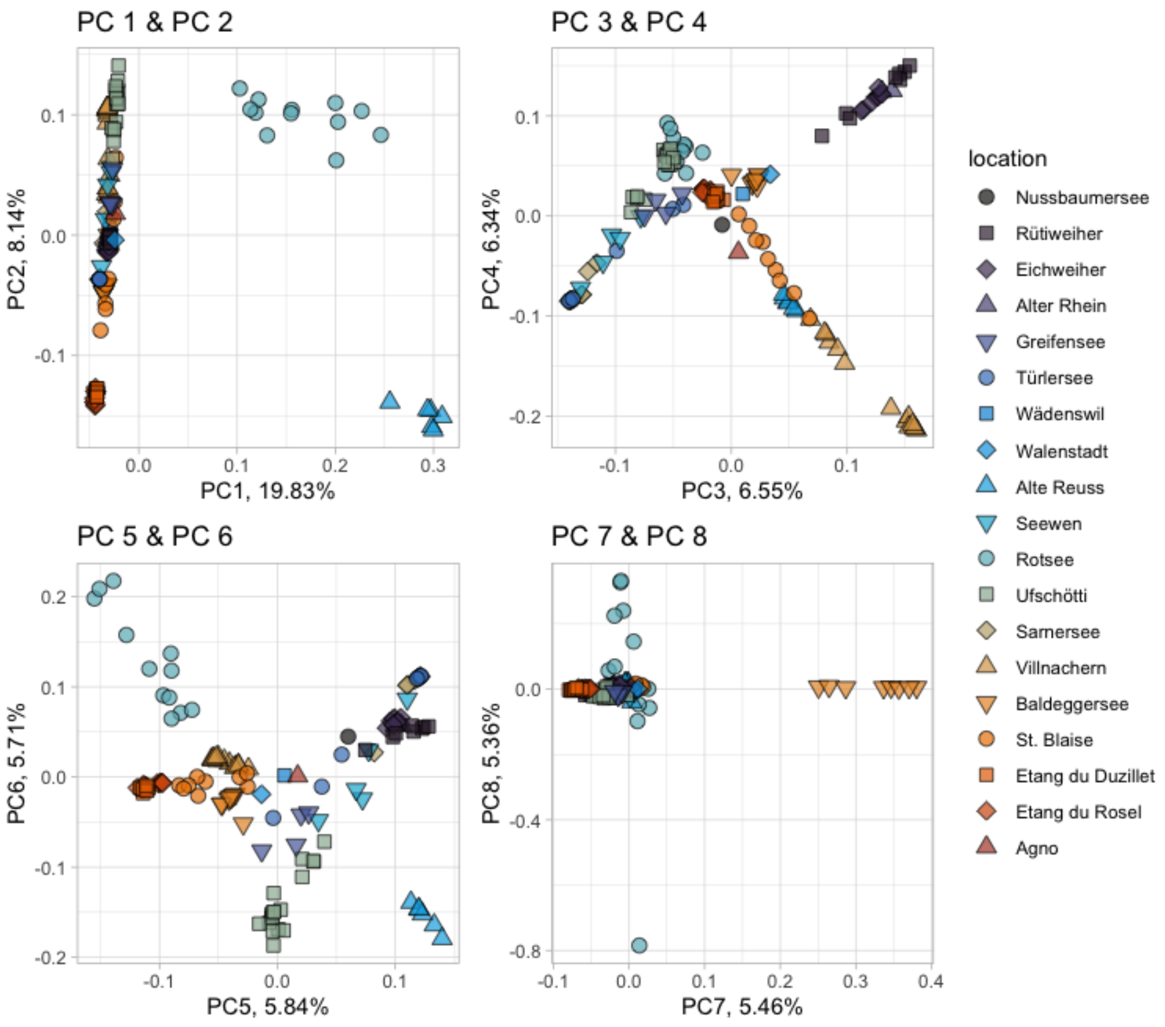

**Figure S13:** Maximum-Likelihood phylogenetic tree of *A. cygnea*, coloured by sampling locality.

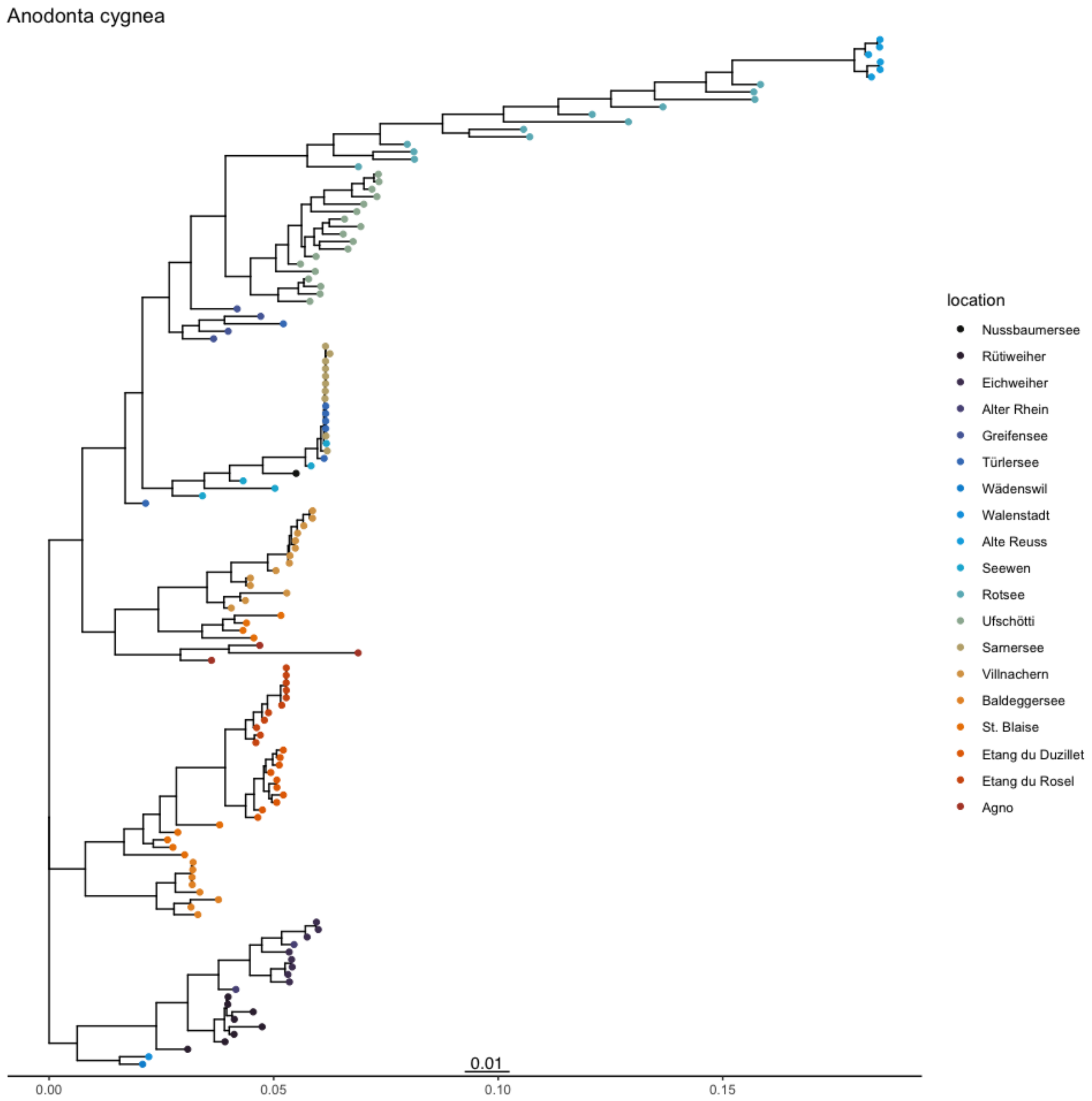

**Figure S14:** Pairwise $F_{ST}$ values among all *A. cygnea* and *A. exulcerata* populations.

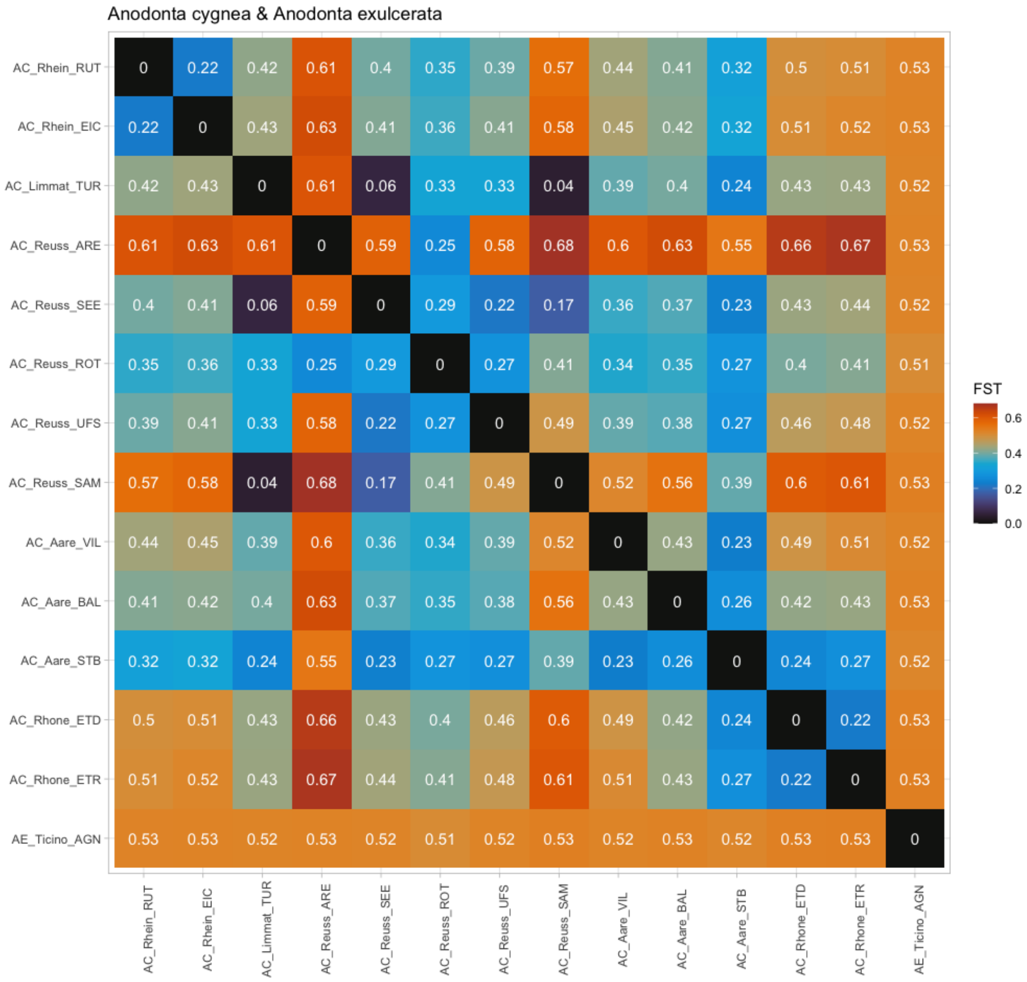

**Figure S15:** Population pairwise average kinship estimates among *A. cygnea* populations.

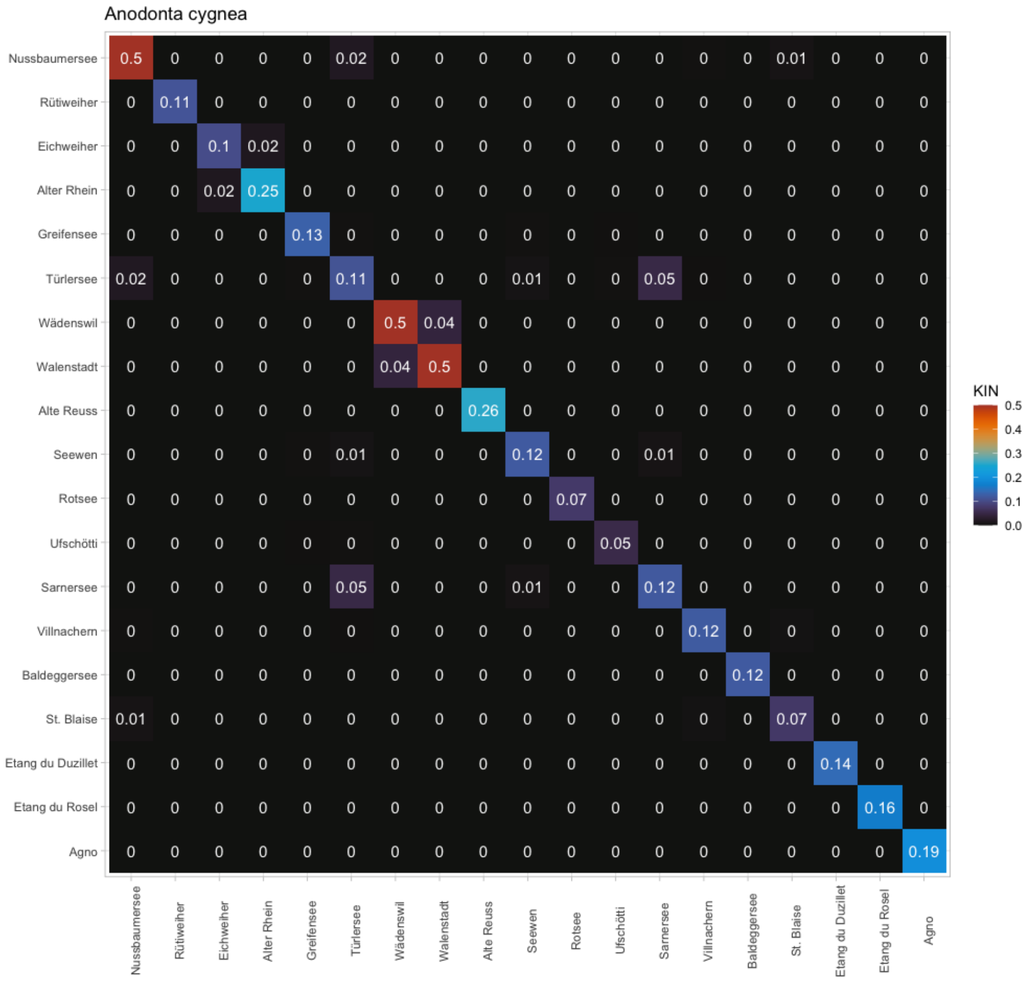

**Figure S16:** Absence of strong association between genetic indicators (observed heterozygosity, inbreeding coefficient, nucleotide diversity, fraction of runs of homozygosity, effective population size) and waterbody size in *A. cygnea.*

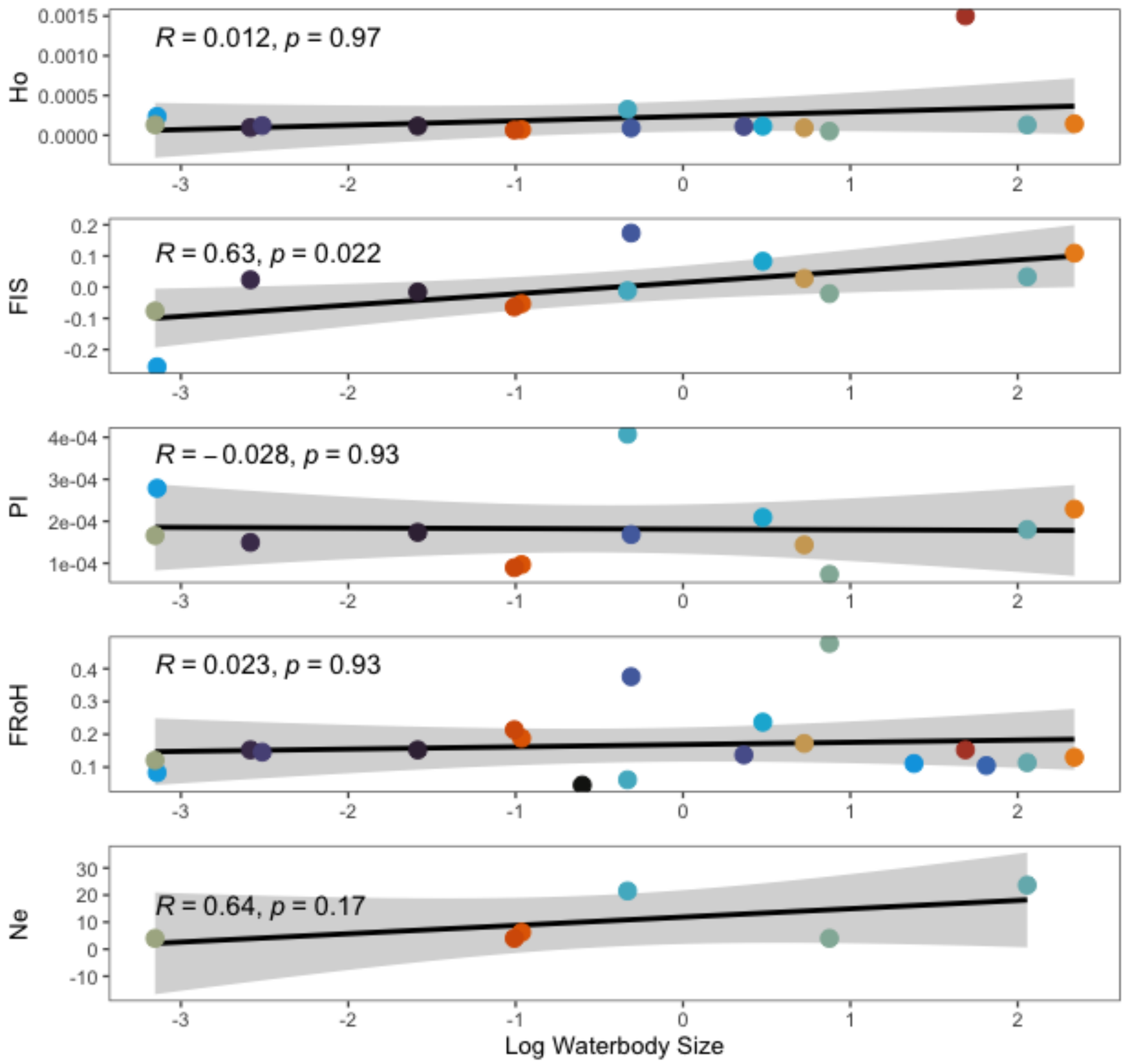

**Figure S17:** Absence of isolation by distance in *A. cygnea.*

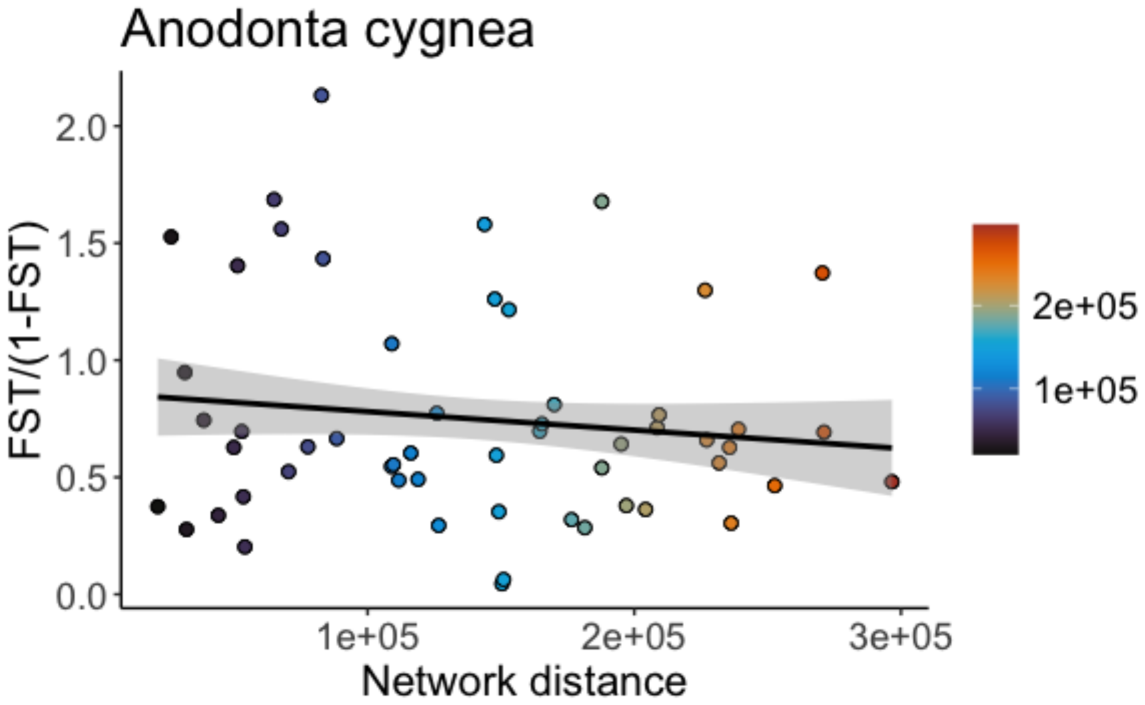
